# Evolutionary specialization of Rad51 and Dmc1 tune recombination outcomes

**DOI:** 10.1101/2025.02.19.639135

**Authors:** Bryan Ferlez, Peter T. Huang, Jingyi Hu, Ilayda Korkmaz, J. Brooks Crickard

## Abstract

Homologous recombination (HR) relies on the RecA-family recombinases Rad51 and Dmc1, which both catalyze DNA strand pairing and exchange but promote distinct recombination outcomes. How these recombinases became functionally specialized remains unclear. Here, we combine phylogenetic analysis with functional assays to define the basis of this specialization. We identify conserved amino acid variation in the DNA-binding Loop 2 region across the archaeal– eukaryotic lineage and show that these residues tune recombinase interactions with double-stranded DNA. Guided by these evolutionary signatures, we engineer Rad51 variants with altered DNA-binding properties and test their functions. Rad51-like variants promote break-induced replication (BIR), whereas substitutions characteristic of Dmc1 suppress BIR and instead favor crossover outcomes, as measured using mitotic crossover assays. These results indicate that evolutionary divergence in Loop 2 alters recombinase binding on reaction intermediates, influencing their stability and progression toward distinct repair outcomes. Thus, evolutionary tuning of recombinase–DNA interactions shift the balance between recombination pathways, providing a mechanistic basis for the functional specialization of Rad51 and Dmc1 during mitotic and meiotic HR.

## Introduction

Homologous recombination (HR) is a conserved pathway that repairs DNA double-strand breaks through template-directed DNA synthesis (Filippo *et al*, 2008; Kowalczykowski, 2015; Symington *et al*, 2014; Szostak *et al*, 1983). In eukaryotes, HR operates in distinct cellular contexts. During vegetative growth, HR promotes the repair of DNA damage and can restart collapsed or damaged replication forks through recombination-dependent pathways such as break-induced replication (BIR) (Heyer, 2007; Kramara *et al*, 2018; Lee *et al*, 2024), whereas during meiosis it repairs programmed DNA breaks and promotes interactions between homologous chromosomes (Arter & Keeney, 2024; Hunter, 2015; Zickler & Kleckner, 2023). These reactions depend on RecA-family recombinases that identify homologous DNA and catalyze strand exchange (Bishop *et al*, 1992; Seitz *et al*, 1998; Shinohara *et al*, 1992; Sung, 1994; West *et al*, 1981). Rad51 functions as the principal recombinase during mitotic HR. Rad51 and the meiosis-specific recombinase Dmc1 cooperate during meiosis, with Dmc1 providing the primary strand-exchange activity (Cloud *et al*, 2012).

Conserved features of all recombinases include the formation of ATP-dependent filaments along DNA (Conway *et al*, 2004; Petassi *et al*, 2026; Shin *et al*, 2025). Productive strand exchange occurs when recombinase filaments assembled on ssDNA search for and engages homologous dsDNA. Initial recognition of short stretches of sequence homology promotes pairing between the invading ssDNA and the complementary strand of the dsDNA, displacing the non-homologous strand to form a displacement loop (D-loop) (De Vlaminck *et al*, 2012; Piazza & Heyer, 2019; Piazza *et al*, 2019; Qi *et al*, 2015; Ragunathan *et al*, 2012; Shibata *et al*, 1980; Yang *et al*, 2020). D-loop formation requires coordinated interactions between two DNA-binding sites within the recombinase filament. DNA-binding site I, formed in part by DNA-binding loops 1 and 2, directly engages the DNA and promotes the strand-exchange reaction (Greenhough *et al*, 2026; Joudeh *et al*, 2025; Luo *et al*, 2025; Yang & Pavletich, 2021; Yang *et al*., 2020), whereas DNA-binding site II stabilizes the displaced strand and prevents its re-annealing. Thus, changes within DNA-binding loops have the potential to influence both the chemistry of strand exchange and the behavior of the recombinase filament.

Despite catalyzing the same fundamental reaction, Rad51 and Dmc1 promote different recombination outcomes, raising the question of how their distinct functions evolved. Rad51 and Dmc1 arose from an ancient gene duplication in the eukaryotic lineage and have retained substantial sequence and structural similarity (Lin *et al*, 2006). Nevertheless, they differ in several biochemical properties, including the kinetics of strand exchange and tolerance for mismatched substrates (Lee *et al*, 2015; Li *et al*, 2021; Steinfeld *et al*, 2019). Additionally, filaments of Dmc1 are more resistant to disruption by regulatory motor proteins such as Rad54, Srs2, and Sgs1 (Bugreev *et al*, 2011; Crickard *et al*, 2018; Crickard *et al*, 2019). These differences could be attributed to DNA binding, but this has not been directly tested. One of the most prominent differences lies within DNA-binding loop 2, which participates in DNA engagement and strand exchange through interactions with the adjacent DNA-binding loop 1 (Luo *et al*, 2021; Xu *et al*, 2021). In Rad51, an aspartic acid within loop 2 forms a conserved salt bridge with an arginine in loop 1, whereas this interaction is disrupted in Dmc1 by substitution of the corresponding residue with glycine or another small amino acid (Luo *et al*., 2021; Xu *et al*., 2021). A second difference occurs at the neighboring position, where Rad51 contains a valine and Dmc1 generally contains proline. These substitutions have been proposed to alter the flexibility of the DNA-binding loops and thereby contribute to differences in strand exchange and mismatch tolerance (Luo *et al*., 2021; Xu *et al*., 2021).

Whether these sequence differences primarily evolved to alter the chemistry of strand exchange, or instead modify the behavior of recombinase filaments, remains unclear. Recombinase filaments must undergo repeated transitions between dsDNA bound and unbound states during HR (Bell & Kowalczykowski, 2016; Haber, 2018; Marin-Gonzalez *et al*, 2025). Their stability can therefore influence multiple stages of the reaction, including homology search, D-loop formation, and subsequent access to the invading DNA end (Hu & Crickard, 2024; Li & Heyer, 2009; Liu *et al*, 2011; Reitz *et al*, 2021; Wright *et al*, 2018). Small changes in filament dynamics could consequently alter recombination outcomes without requiring changes to the core chemistry of strand exchange. Determining whether such dynamic properties were themselves targets of evolutionary specialization requires connecting sequence variation across the recombinase family to its effects on recombination in cells.

Here, we investigated the evolutionary history and functional consequences of variation within Rad51 DNA-binding loop 2. We first examined recombinase sequences across Archaea and Eukarya and reconstructed the ancestral state of the Rad51/Dmc1 lineage. These analyses identified a Pro-Asp loop configuration as an ancestral state that predates the divergence of Rad51 and Dmc1, whereas the Val-Asp configuration characteristic of Rad51 arose later in the Rad51 lineage. We then introduced ancestral and Dmc1-like substitutions into yeast Rad51 and examined their effects on recombination outcomes, D-loop formation and extension, ATP hydrolysis, and filament dissociation. These experiments reveal that changes within a small region of the DNA-binding loop can alter recombinase dynamics and, in turn, shift the balance among recombination outcomes. Our findings support a model in which evolutionary divergence of Rad51 and Dmc1 involved modification of the intrinsic DNA binding of an ancestral recombinase, linking sequence evolution within a DNA-binding loop to the regulation of recombination outcomes.

## Results

### Analysis of recombinase diversity Across the Archaea/Eukarya Tree

An existing hypothesis is that Dmc1 was duplicated from an existing Rad51 gene and gained a novel function, becoming useful for meiosis. If this were true, then we would expect lineage-specific amino acids that contribute to this novel function to only exist after the split between Rad51 and Dmc1 in eukaryotes. Lineage-specific amino acids that are thought to differentiate Rad51 and Dmc1 function occur in two DNA binding loops, L1 and L2. These loops represent a point of collaboration between two adjacent protomers in all domains of life (Bell & Kowalczykowski, 2016). High-resolution structures demonstrate that an arginine in L1 (R235 in *H. sapiens* RAD51 and R293 in *Saccharomyces cerevisiae* Rad51) forms a salt bridge with an L2 aspartic acid (D274 in *Homo sapiens* RAD51 and D332 in *S. cerevisiae* Rad51) from an adjacent protomer (**Fig EV1A**). This interaction is conserved in Rad51. However, in Dmc1, this interaction is lost because the corresponding aspartic acid is replaced by a glycine or another small amino acid such as Serine or alanine (**Fig EV1A**). A second, less appreciated difference involves a hydrophobic contact between the two loops. A Leucine residue in L1, which is completely conserved in eukaryotic recombinases and highly conserved in archaea, forms a specific hydrophobic interaction with a Valine in L2 of Rad51 (V273 in *H. sapiens* RAD51 and V331 in *S. cerevisiae* Rad51). In Dmc1, the L1 Leucine is retained, whereas the corresponding L2 residue is a proline (P274 in *H. sapiens* DMC1 and P223 in *S. cerevisiae* Dmc1), likely disrupting this interaction. Therefore, Rad51 and Dmc1 differ in both the salt bridge and hydrophobic contact that connect L1 and L2, potentially altering how these loops are stabilized during strand invasion.

To determine if loss of the salt bridge existed prior to the differentiation of the ancestral eukaryotic recombinase, we obtained recombinase sequences from Archaea by performing BLASTp with *S. cerevisiae* Rad51 (Uniprot P25454), *H. sapiens* RAD51 (Uniprot Q06609), and *S. solfataricus* RadA (Uniprot Q55075) sequences as queries to Archaea. We initially queried each of the four major archaeal groups, divided into the TACK superphylum (Thaumarchaeota, Aigarchaeota, Crenarchaeota, and Korarchaeota), Euryarchaeota (EURY), the DPANN superphylum (Diapherotrites, Parvarchaeota, Aenigmarchaeota, Nanoarchaeota, and Nanohaloarchaeota), and the Asgard superphylum. We returned 758, 782, 621, and 198 high-confidence RadA protein sequences from these four major groups, respectively (**Fig EV2A and Extended Data 1**) (Nguyen *et al*, 2014). Many of these DNA sequences were generated from metagenomic data. Because of this we were careful in which sequences we scored as actual recombinase enzymes. If sequences lacked the Walker A or Walker B motifs they were excluded from the analysis. Likewise, if there was no obvious DNA binding loops the sequence was removed from the analysis. After generating protein sequence alignments for each of these groups, we generated unrooted maximum-likelihood phylogenetic trees (**Fig EV3A and Extended Data 1**).

We used these trees to generate a larger phylogenetic tree that included an additional 43 eukaryotic Rad51 sequences and 28 eukaryotic Dmc1 sequences, as well as 81 bacterial sequences comprising *Escherichia coli* RecA, and two sequences selected from each of 40 eubacterial phyla. The addition of these sequences generated a large, unrooted maximum-likelihood phylogenetic tree composed of 2511 sequences. The clustering of TACK, EURY, and DPANN sequences within groups reflecting their representative superphyla was primarily supported by SH-aLRT and UFBS statistical tests (**Fig 1A**) (Hoang *et al*, 2017; Nguyen *et al*., 2014) As expected, the Asgard Archaea grouped closest to the eukaryotic proteins.

**Figure 1:**
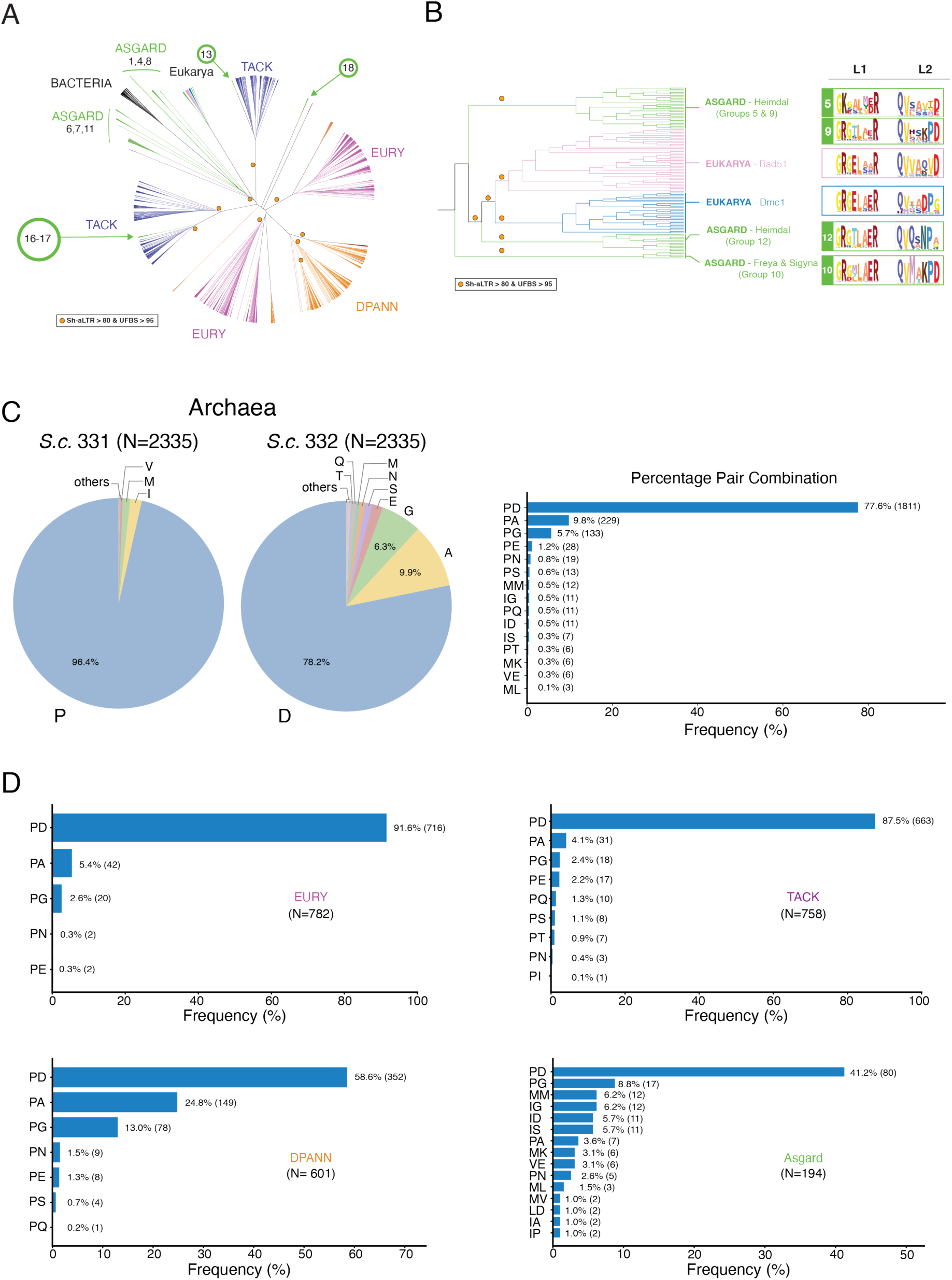
Dmc1 binding loop is common in Archaea. **(A).** Cladogram view of maximum likelihood phylogenetic tree built from recombinase sequences spanning the domains of Archaea (TACK (Purple), Euryarchaeota EURY (Magenta), DPANN (Orange), and Asgard (Green)), Eubacteria, and Eukarya. The orange dots represent statistical significance using SH-aLRT and UFBS (SH-aLRT >80 & UFBS >95). **(B).** Cladogram view of the phylogenetic tree with an expanded view of the split between Archaeal and Eukaryotic recombinases. To the left is a sequence logo for the DNA-binding loops within each clade. The yellow dots represent statistical confidence in the split of the branches. **(C).** Single-position amino acid distribution and paired-residue frequencies across the combined archaeal dataset, comprising Asgard, TACK, EURY, and DPANN sequences. Pie charts show amino acid frequencies at positions equivalent to *S. cerevisiae* Rad51 residues 331 and 332, analyzed independently at each position. For each position, *N* indicates the number of sequences containing a non-gap residue. The horizontal bar plot shows the 15 most frequent two-residue combinations, calculated among sequences containing non-gap residues at both positions. Values beside each bar indicate the percentage of valid pairs, followed by the corresponding sequence count in parentheses. Less frequent pair combinations not displayed in the bar plots remained included in the denominator. **(D).** Frequencies of paired-residue combinations analyzed separately for EURY, TACK, DPANN, and Asgard archaea. For each plot, *N*indicates the number of sequences containing non-gap amino acids at both positions.

To determine which group of Asgard Archaea was closest to the last common eukaryotic ancestor, we used statistical confidence in the branching of the Asgard recombinase tree to define eighteen distinct groups of recombinases (**Fig EV3B and EV3C**). Our stringent criteria for accepting a protein as a recombinase had identified that four of these groups, 2, 3, 14, and 15 failed the established criteria check and had been removed from the analysis. Based on this analysis, the nearest relative of Rad51/Dmc1 in Archaea was group 12, which was composed entirely of the Heimdall group of Asgard Archaea (**Fig 1B**). This is consistent with previous observations that the Heimdall group is a sister lineage of modern eukaryotes (Eme *et al*, 2023; Liu *et al*, 2021; Spang *et al*, 2018; Zaremba-Niedzwiedzka *et al*, 2017). When we evaluated the L1 and L2 sequences in Asgard group 12, we were surprised to find that they all had lost the salt-bridge between L1 and L2 and had a proline in place of the valine present in Rad51.

We next asked how common the Dmc1-like loop type was in Archaea. Proline was present in 96.4% of Archaeal RadA, and aspartic acid was present in 78.2% of sequences (**Fig 1C**). The proline- aspartic Acid pair (-PD) was the most common loop type in Archaea (77.2%) (**Fig 1C and Fig EV4**). This is likely a hybrid of Rad51 and Dmc1, which retains the salt bridge but has a modified hydrophobic contact. The second highest combination was Proline-Glycine and Proline-Alanine at 9.8 and 5.7%, respectively. This data showed that in around 15-16% of RadA sequences in Archaea there is a loss of the salt bridge and an alteration to the hydrophobic contact, like the Dmc1 configuration observed in eukaryotes. Valine was found in only 0.2% of sequences in Archaea. This loop variation could be analogous to Rad51 however it was one of the rarest variants in Archaea.

We evaluated whether the distribution of these amino acids was uniform across major archaeal clades. Proline was present at the first position 100% of the time in EURY, TACK and DPANN (**Fig 1D**). In EURY and TACK, this proline was accompanied by an aspartic acid in 90% of cases, whereas a minor fraction (6-8%) carried Dmc1-like PG or PA combinations. valine at this position was not observed in these major clades. In DPANN only 58.6% of sequences were a -PD combination, and 37% were -PA/PG, marking this superphylum with the highest level of Dmc1- like loops (**Fig 1D**). Again, valine was not observed in this superphyla. As a group the DPANN archaea have small genomes and exhibit higher levels of horizontal gene transfer. While there is no direct evidence that RadA from this group behaves more like Dmc1, the correlation between loop type and higher levels of genetic exchange is notable and may represent an evolutionary strategy to increase genetic exchange in these organisms. The Asgard group had much higher sequence diversity. However, the two most common amino acid pairs were still -PD (41%) and - PG (8.8%). There are additional variations at these positions, but valine was still uncommon (3.1%) and occurred only in combination with glutamic acid (**Fig 1D**). We cannot rule out the possibility that the high level of variation in Asgard Archaea is due to larger levels of metagenomic protein sequences used in this analysis. However, from this data we can conclude that the two most prevalent amino acids in these positions are -PD, representing the common type in Archaea, and - PA/-PG, representing Dmc1-like loops. We can also conclude that Rad51-like loops are uncommon and may be non-existent in Archaea.

### Molecular reconstruction

Based on our phylogenetic analysis we hypothesized that the last common ancestor of Rad51 and Dmc1 (LEADR) likely had DNA binding loops that resembled a common Archaeal organization (-PD). However, we could not rule out that the ancestor may have a Dmc1 organization (-PG). To differentiate between these two possibilities, we used an ancestral reconstruction method to predict the ancestral state of both Rad51 and Dmc1. More detailed descriptions of our methodology in reconstructing the ancestor can be found in the Materials and Methods. Here we summarize several critical steps and our findings. To perform Ancestral Sequence Reconstruction (ASR), we used an existing data set (Matsuo *et al*, 2025), supplemented with 2 additional Rad51 sequences, to infer a maximum-likelihood phylogeny of RadA, Rad51 and Dmc1 using IQ-TREE (Wong *et al*, 2026). The resulting tree was then rerooted on the branch separating archaeal RadA from the eukaryotic Rad51/Dmc1 clade using custom code (Nguyen *et al*., 2014). From the rerooted tree, we identified several ancestral nodes for sequence reconstruction, including the common ancestor of archaeal RadA and eukaryotic recombinases (Anc_RadA_Euk), the ancestral eukaryotic recombinase shared by Rad51 and Dmc1 (LEADR; Anc_Euk), and the respective ancestors of the Rad51 and Dmc1 clades (Anc_Rad51 and Anc_Dmc1) (**Fig 2A**). The sequences reconstructed at these internal nodes should be viewed as statistical inferences of ancestral states rather than directly observed sequences.

**Figure 2:**
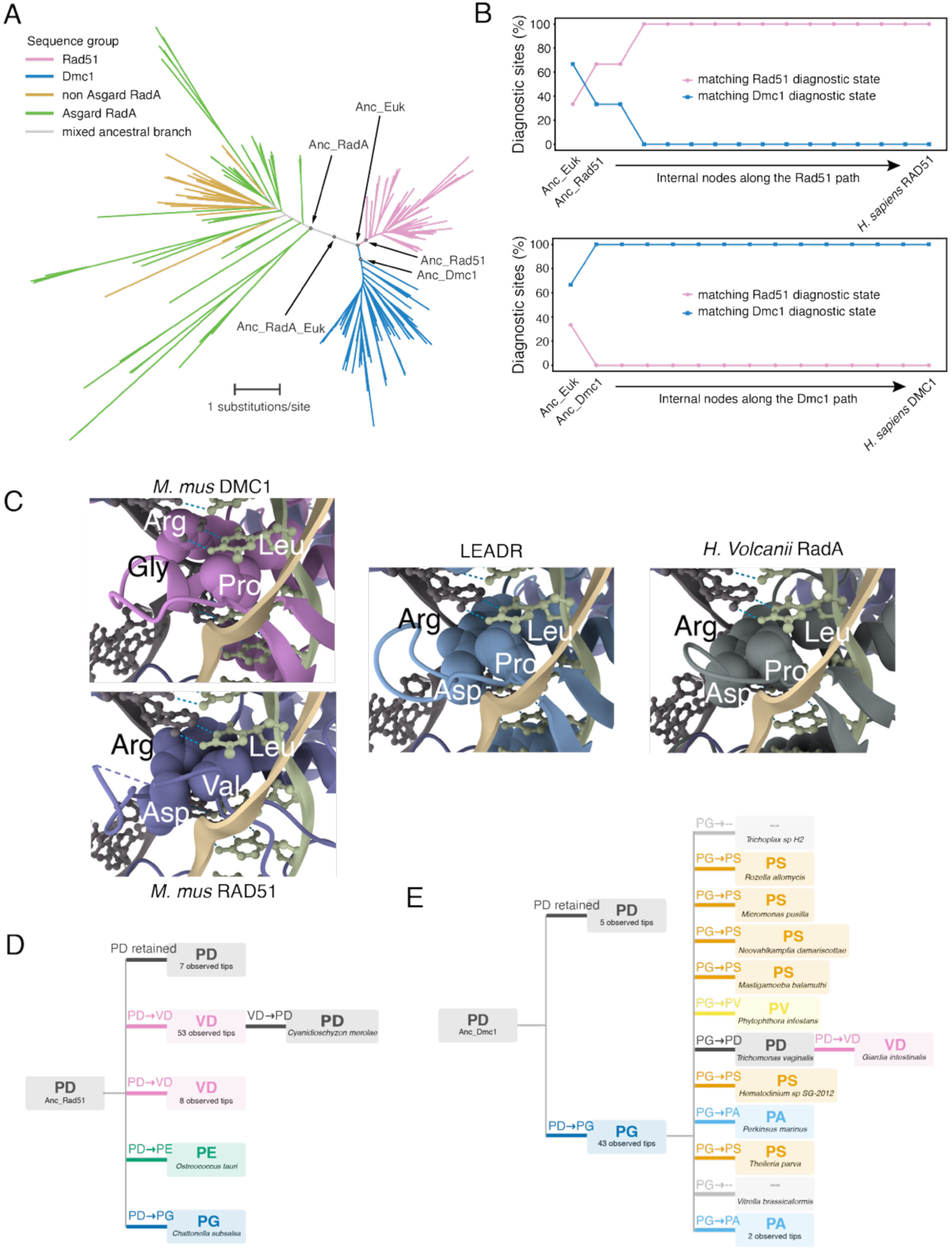
Reconstruction of the evolution of eukaryotic recombinase loop 2. **(A)**. Maximum-likelihood phylogeny of archaeal RadA and eukaryotic Rad51 and Dmc1 sequences. Branches are color according to the sequence group, and gray branches represent the ancestral region connecting archaeal and eukaryotic recombinases. Ancestral nodes reconstructed in this study are indicated. The scale bar represents substitutions per site. **(B)**. Evolutionary trajectories of Rad51- and Dmc1-diagonostic sites along the lineages leading from Anc_Euk to *H. sapiens* RAD51 (top) and *H. sapiens* DMC1 (bottom). The percentage of diagnostic sites matching the Rad51-type or Dmc1-type states are shown for successive internal nodes along each path. **(C)**. Structural comparison of the Loops 1 and 2 in *Mus musculus* DMC1, *M. musculus* RAD51, LEADR, and *H. volcanii* RadA. LEADR denotes the recommended ancestral eukaryotic recombinase reconstructed from the curated RadA/Rad51/Dmc1 dataset. Structures of DMC1, LEADR and RadA were predicted using AlphaFold3 and structurally aligned to the experimentally determined mouse RAD51-D-loop complex (PDBID: 9UI8). Close-up views show the positions of loops 1 and 2 relative to the D-loop DNA, with focal residues shown as spheres and labeled. **(D).** Simplified map of amino acid pair-state evolution at positions equivalent to *S. cerevisiae* Rad51 residues 331–332 from the reconstructed Anc_Rad51 state. The ancestral PD state was retained or independently changed to VD, PE, or PG, with a secondary VD-to-PD transition. Numbers indicate the observed tips represented by each branch, and species names are shown for singleton events. Independent transitions resulting in the same pair state are displayed separately. A detailed phylogenetic representation of these transitions is provided in Supplementary Fig. 9. **(E).** Simplified map of amino acid pair-state evolution from the reconstructed Anc_Dmc1 state. The ancestral PD state was retained or changed to PG, followed by multiple lineage-specific secondary transitions from the PG state. Numbers indicate the observed tips represented by each branch, and species names are shown for singleton events. Independent transitions resulting in the same pair state are displayed separately.

To generate a robust proteins sequence for LEADR, we analyzed two unique datasets with different multiple-sequence alignment methods (MUSCLE or MAFFT) (Edgar, 2022). For each alignment we considered protein alignments untrimmed or trimmed with trimAl (Capella-Gutiérrez *et al*, 2009), creating eight separate analysis pipelines. In each resulting phylogeny, we identified the same topologically defined ancestral eukaryotic recombinase node (LEADR), which separated the Rad51 and Dmc1 clades from the RadA lineage. The consistent recovery of this node across all eight analyses allowed us to compare the reconstructed LEADR sequences among pipelines.

The full-length ancestral reconstruction converged on a 341-aa sequence. Among the amino acid positions that could be directly compared across all eight ASR pipelines (201 positions in total), 179 (89.1%) had the same maximum-posterior ancestral state in every run, and 163 (81.1%) reconstructed the same residue with posterior probability ≥ 0.80 in all eight analyses. Only 22 positions showed pipeline-dependent maximum-posterior states. The remaining 140 positions were not represented in all eight analyses, largely because some alignment columns were removed during trimming rather than because of disagreement over the ancestral residue presence. Importantly, 138 of these 140 positions were retained in all four untrimmed full-length reconstructions, indicating that their inclusion in the ancestral sequence was also highly consistent across independent alignments. For the recommended full-length sequence, positions from residue 34 onward were assigned by taking, at each position, the amino acid with the highest unweighted mean posterior probability across the pipelines in which that position was present. The principal remaining uncertainties involved the alignment and exact composition of the acidic, low-complexity 33-aa N-terminal region, the retention of the position corresponding to human RAD51 H275, and several core positions with closely competing ancestral states (**Fig EV5A**). Only two positions showed sensitivity in residue presence among the four untrimmed reconstructions. The four untrimmed full-length candidates shared >90% pairwise sequence identity (**Fig EV5B**).

To determine if the ancestor was generally more Dmc1-like or Rad51-like we evaluated several diagnostic residues. These positions differ between Dmc1 and Rad51, are present in >70% of sequences in each lineage, and show >80% conservation of the lineage-specific amino acid among homologs in which the position is present (**Extended Table 1**). This analysis suggested that the ancestor of Dmc1 and Rad51 had more diagnostic residues in common with Dmc1 than Rad51, and indicates the ancestor was more like Dmc1 from this perspective (**Fig 2B and Fig EV6AB and Fig EV7AB**). However, in total sequence composition the ancestor was more like Rad51(**Fig EV6 and Fig EV8)**.. In summary, our data suggests that the ancestor is a mixture of the two recombinases.

All models generated from ASR contained a salt bridge between L1 (Arginine at equivalent R235 in *H. sapiens* RAD51) and L2 (Aspartic acid at equivalent D274 in *H. sapiens* RAD51) and had the -PD configuration of L2. The presence of this configuration implies the loss of the salt bridge evolved in Dmc1 lineage and the gain of valine in L2 in the Rad51 lineage. We used Alphafold to generate structural models of LEADR with ssDNA and with a homologous dsDNA donor (**Fig 2C**). We aligned this prediction with existing structures of Rad51 during D-loop formation (**Fig 2C**) (Luo *et al*., 2025). The alignment on the existing Rad51 structure allowed us to observe predicted changes in the L1-L2 interaction during strand pairing. LEADR exhibited a gap in L1- L2 interaction because the proline is predicted to interact less efficiently with leucine. This interaction was more like those observed in Archaea (*Haloferax volcanii*), and it was also distinct from that observed for both Rad51 and Dmc1 (**Fig 2C**).

The reconstruction also allowed us to follow the evolution of L2 configuration within the phylogenetic tree. A transition was identified along a branch when the inferred L2 configuration differed between a parent node and its immediate descendant node or terminal tip. The most common conversion in Rad51 was from proline-aspartic acid to valine-aspartic acid (**Fig 2D and Fig EV9**). Consistent with the ancestral reconstruction, the inferred ancestral -PD configuration was retained in red and green algal lineages that diverged relatively early in eukaryotic evolution. Dmc1 had a significantly larger number of transitions within the tree. However, except for in *three examples,* a proline was maintained at the first position, and most of the variation occurred at the second position in the L2 (**Fig 2E and Fig EV10**). From this we conclude that Rad51 and Dmc1 functionally diverged from LEADR by variations in the interactions between L1 and L2, and that the original L2 configuration was -PD.

### Substitution of Dmc1-like loops in Rad51-like results in Crossover outcomes

Our analysis of recombinase evolution is consistent with a hypothesis that the last common ancestor of Rad51 and Dmc1 had DNA binding loops that were a mixture of the two recombinases. Therefore, the DNA binding site of Rad51 has undergone evolutionary changes that have made this recombinase more efficient at double strand break repair during mitosis. Similarly, the DNA binding site of Dmc1 underwent evolutionary changes that enhanced its specialization for meiotic recombination, promoting homolog bias and crossover formation. The changes in Rad51 lineage are likely linked to the evolution of a valine residue within L2. We hypothesized that replacing proline with valine could alter the L1-L2 interface by removing the conformational constraint imposed by proline while introducing a larger hydrophobic side chain that may favor interaction with the leucine residue in L1. This model predicts that further increase in the hydrophobic side-chain surface at this position could enhance this interaction. We therefore substituted phenylalanine for valine in L2 to test whether this would further increase strand-exchange efficiency.

Therefore, we made a series of mutations in L2 to examine the effects on the recombination reaction *in vivo*. This included *rad51-V331F* (*rad51-FD*), *RAD51* (*RAD51-VD*), *rad51-V331P* (*rad51-PD*), and *rad51-V331P, D332G* (*rad51-PG*) (**Fig 3A**). We initially performed experiments in an *S. cerevisiae* strain in which the expression of the HO endonuclease could induce a double strand break cutting a site within a *MATa* gene on chromosome V. A template to repair the break is located on chromosome III at the *MATa* locus. The repair template is an incomplete version of the *MATa* gene, and the use of this substrate leads to gene conversion outcomes (Ira *et al*, 2003). Failure to undergo gene conversion results in cell death and failure to grow on galactose-containing media (**Fig 3B**).

**Figure 3:**
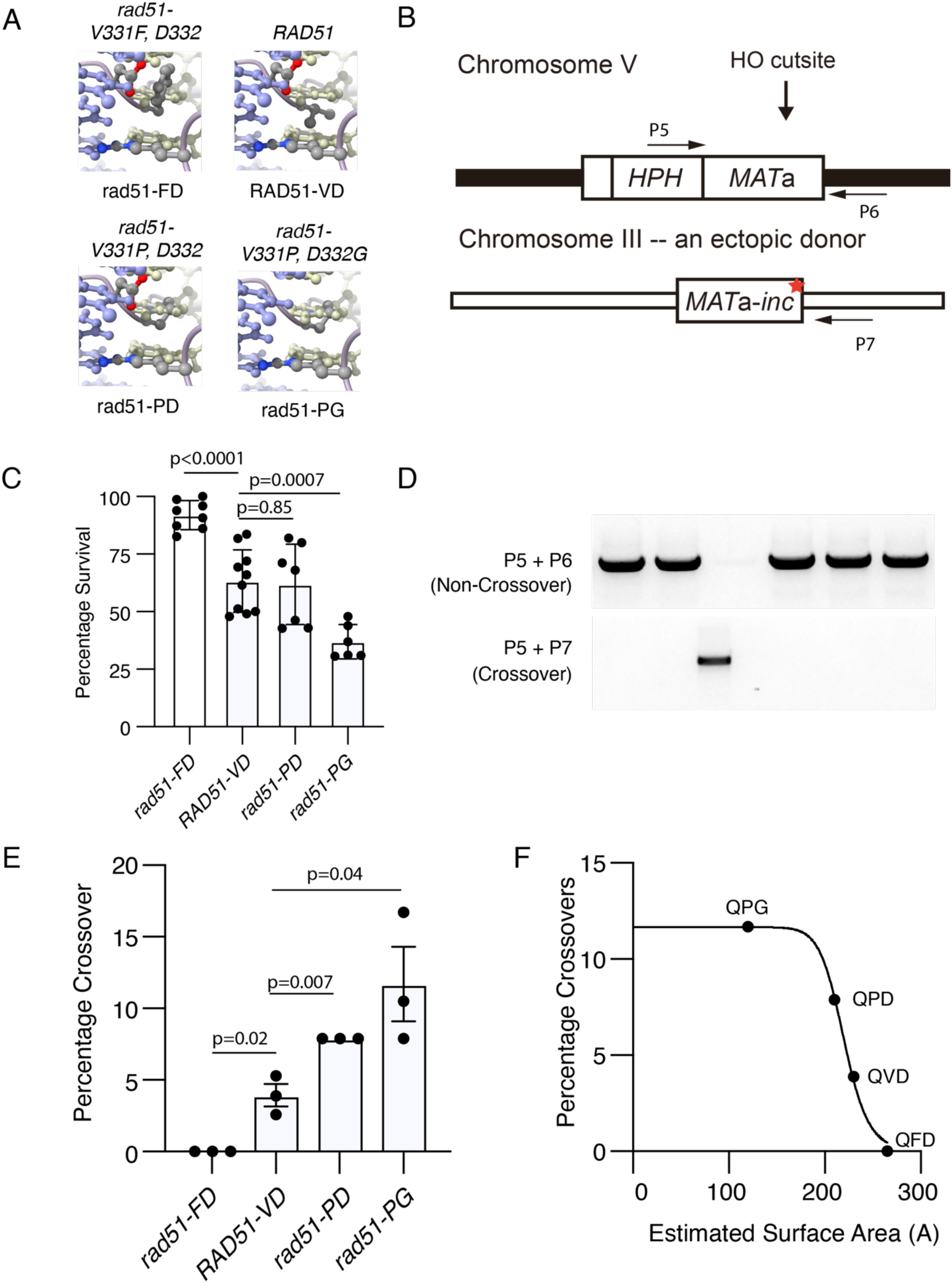
Substitutions in DNA binding loop 2 can promote genetic exchanges. **(A).** Images representing the amino acid substitutions made in the DNA-binding loop 2 of Rad51. The images were generated by substituting the amino acids into an existing Rad51 structure. **(B).** Schematic diagram illustrating the design of the double strand break repair reporter used to test the effects of loop substitutions. An HO endonuclease cut site was located on chromosome V and could be repaired from a template on chromosome III. **(C).** Graph representing the survival percentage of *rad51-FD, RAD51-VD, rad51-PD,* and *rad51-PG*. The bar represents the mean, and the error bars represent the standard deviation of at least four independent experiments. **(D).** Representative gel image illustrating PCR product using primer pairs to detect non-crossovers (P5 and P6) and primer pairs to detect crossovers (P5 and P7). **(E).** Graph representing the percentage of PCR products that occurred using crossover primers for *rad51-FD, RAD51-VD, rad51-PD,* and *rad51-PG.* The bars represent the mean, and the error bars the standard deviation for three independent experiments. **(F)**. Graph comparing the estimated surface area for QFD, QVD, QPD, and QPG in DNA binding loop 2 as a function of crossover percentage.

In our hands, *RAD51-VD* promotes gene conversion and survival ∼60-65% of the time (**Fig 3C**). Survival was significantly increased in the *rad51-FD* mutant to around 85%. The ancestral *rad51- PD* had the same survival rate as WT, and *rad51-PG* presented a 2-fold defect in survival (**Fig 3C**). From this, we conclude that mutation of both the salt bridge and valine 331 resulted in an alteration to survival during ectopic repair. The repair from an ectopic site allows the assessment of further genetic exchange between chromosome III and chromosome V. We hypothesized that the *rad51-PG* mutant would be more Dmc1-like and would have an increase in mitotic crossovers.

Whether this occurs in a surviving colony can be determined by using specific pairs of PCR primers (**Fig 3B D**). We screened approximately 140 colonies per mutant from three separate plates and quantified the percentage of colonies that exhibited crossover outcomes. In *RAD51-VD,* this was approximately 3-5%, as previously reported (Ira *et al*., 2003). Mutation of valine 331 to proline resulted in significant increases in crossover outcomes, with the *rad51-PD* increasing to 7.9% and the *rad51-PG* increasing to 11.7% (**Fig 3E**). The conversion of valine 331 to phenylalanine resulted in a reduction in crossover. We tested a larger number of colonies for the *rad51-FD* mutant and were unable to observe a crossover outcome. It could be that we did not sample enough colonies, and while the measurement is zero, all we can say is that the percentage of crossover outcomes is less than 0.75% (**Fig 3E**).

In all cases, changes in crossover percentages were impacted by the surface area of the residue at position 331. We observed that this data appeared to follow a trend line, and we calculated the combined theoretical surface accessible regions for the glutamine (Q330), valine (V331), and aspartic acid (D332) and variations of these three amino acids. In good agreement with our observations, the various Rad51 mutants illustrated a decrease in crossover outcomes as the surface area of these three amino acids increased (**Fig 3F**). These data suggest that the identity of the amino acids at 331 and 332 in Rad51 impacts the probability of genetic exchange during recombination, and this site can be modified during evolution. Importantly, it suggests that Rad51 has evolved to be less efficient at genetic exchange.

### D-loop capture and extension

The overall efficiency of the recombination reaction is regulated by how quickly recombinase filaments find a homologous template to facilitate repair, and how effectively they are removed from the dsDNA to allow DNA replication to repair the break (Li & Heyer, 2009). We hypothesized that the differences observed in survival and crossover outcomes may be due to the efficiency of the recombinase’s on-off dynamic. To measure this, we used a previously established assay to test the efficiency of intermediate step formation in the HR reaction. The assay uses HO endonuclease to create a DSB in the genome. As in the ectopic repair assay, the repair template is located on a different chromosome (**Fig 4A**). Location of the homologous site results in D-loop formation, and this intermediate can be trapped by psoralen crosslinking. By isolating the crosslinked intermediate the efficiency of D-loop formation can be measured by qPCR. This approach has previously been used to measure the formation and stability of D-loop intermediates (Hu *et al*, 2026; Piazza *et al*., 2019; Reitz *et al*, 2022). This assay lacks a second end for capture, and unlike the gene conversion experiments used earlier, DNA repair cannot go to completion. The readout is strictly the formation of recombination intermediates and the rate at which they occur (**Fig 4A B**).

**Figure 4:**
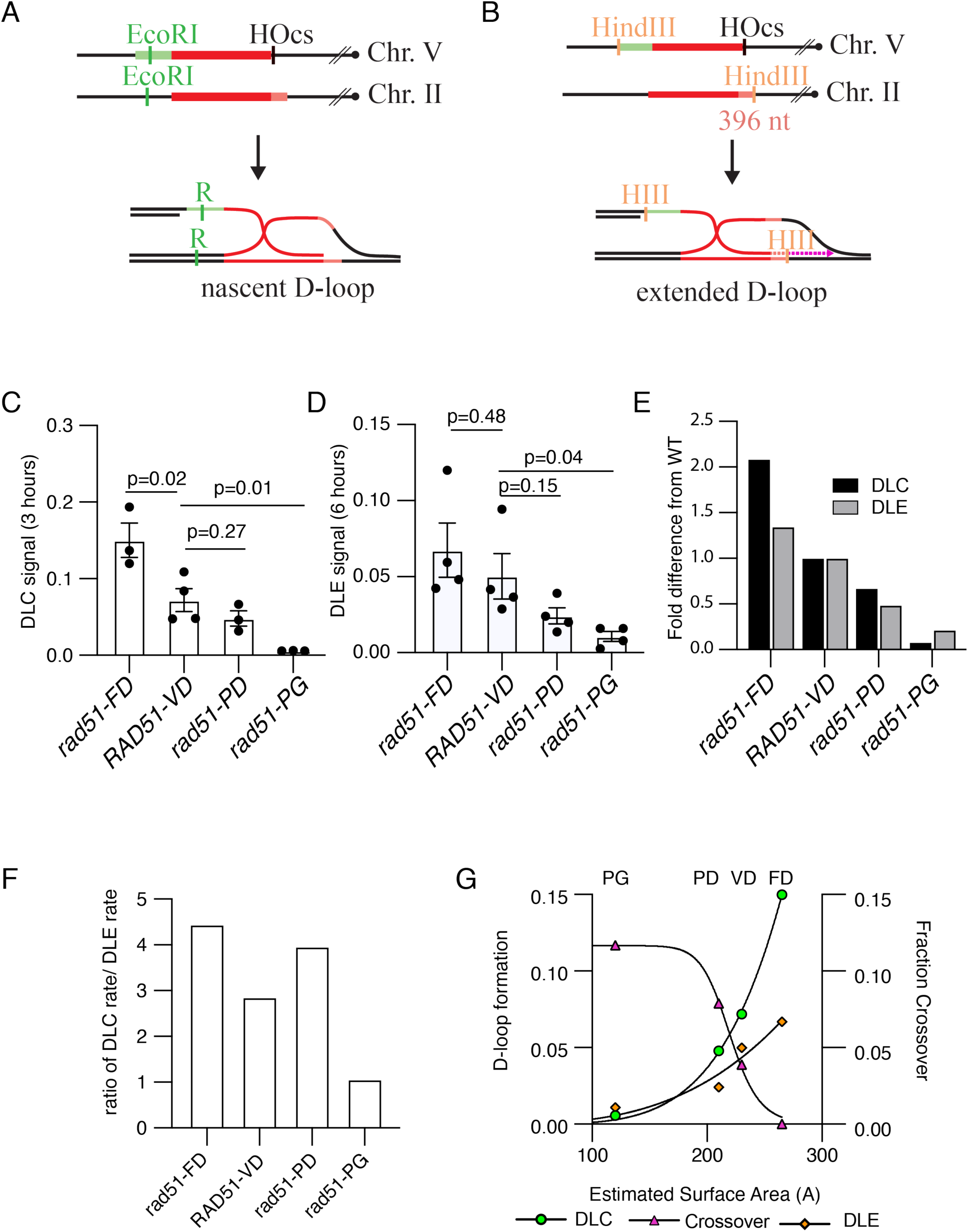
DNA binding loop 2 controls intermediate formation *in vivo*. **(A).** Schematic diagram illustrating the substrate for the in vivo D-loop capture (DLC) assay. **(B).** Schematic diagram illustrating D-loop extension (DLE) assay. **(C).** Graph representing results for the DLC assay at 3 hours for *rad51-FD, RAD51-VD, rad51-PD*, and *rad51-PG*. The bar represents the mean, and the error bars represent the standard deviation for at least three independent experiments. (D). Graph representing results for the DLE assay at 6 hours for *rad51-FD, RAD51- VD, rad51-PD*, and *rad51-PG*. The bar represents the mean, and the error bars represent the standard deviation for at least three independent experiments. **(E).** Graph representing the fold difference from RAD51-VD for the DLC (Black) and DLE (Gray) assay. **(F).** Graph representing the ratio of the DLC rate by the DLE rate for *rad51-FD, RAD51-VD, rad51-PD*, and *rad51-PG*. (G). Graph comparing the efficiency of D-loop formation to the percentage of crossovers that form within a population. This is plotted against the estimated surface area for QFD, QVD, QPD, and QPG in DNA binding loop 2.

Based on the survival observed in the gene conversion assay, we reasoned that the formation of D-loops would be lower or delayed in the *rad51-PG* mutation and would be enhanced in the *rad51- FD* mutant. This would reflect a faster D-loop formation for the *rad51-FD* mutant and a slower D-loop formation for the *rad51-PG* mutant. When we evaluated D-loop capture efficiency, there was a significant two-fold increase in the D-loop capture for the *rad51-FD* allele (**Fig 4C**), and a 10-fold defect in the *rad51-PG* mutant (**Fig 4C**). The *rad51-PD* allele did not show a significant defect in D-loop capture. However, it was lower than wild type and fit an expected trend (discussed below).

These strains can also be used to measure D-loop extension (**Fig 4B**). This measures DNA extension by quantifying D-loops that have extended the recipient DNA by at least 396 nucleotides. Both the accessibility of the 3’ end of the DNA and the extension of DNA polymerase from the D-loop are regulated by recombinase occupancy (Li & Heyer, 2009). Therefore, the overall extension is correlated with the efficiency of recombinase removal. Our measurements of D-loop extension in the *rad51-FD* strain did not result in an increase in extension efficiency higher than wild type (**Fig 4D**), suggesting this mutant may decouple D-loop formation from extension. Likewise, the *rad51-PD* mutant was lower than the wild type. The *rad51-PG* was 5-fold lower than wild type, suggesting that this mutant was less deficient in allowing extension than in promoting capture (**Fig 4D**).

For our experiments, the DLC and DLE samples were collected from the same culture and are an extension of the same experiment, allowing comparison of capture products and extension products. We measured the fold differences from WT and compared the two different readouts, extension or capture. From this analysis, we can see that the fold difference in capture is higher than the fold difference in extension for both *rad51-FD* and *rad51-PD*. In contrast, the fold difference for the *rad51-PG is* higher in extension (**Fig 4E**), suggesting these mutants may impact different steps in D-loop metabolism.

Comparing the rates of the different Rad51 mutants also allowed us to monitor the progress of the total reaction. To measure this, we divided the D-loop capture or D-loop extension products by the time at which the sample was collected and then took a ratio of the rates. This analysis showed that both the *rad51-FD* and *rad51-PD* mutants had higher ratios than the wild type. This could be caused by an increase in capture efficiency (*rad51-FD*), or a decrease in extension efficiency (*rad51-PD*) (**Fig 4F**). In contrast, the *rad51-PG* mutant had a lower ratio than the wild type, suggesting a stronger defect in D-loop capture.

As with the fraction of crossover outcomes, we plotted the D-loop capture and extension efficiency as a function of amino acid surface area. We found the data fit a trend where more surface area in the binding loop led to more efficient D-loop formation. This trend was the inverse of the crossover outcomes, suggesting that crossover efficiency was related to the rate of D-loop formation (**Fig 4G**). From this analysis, we conclude that the rate of D-loop formation likely impacts the amount of crossover formation and impacts survival in the gene conversion assay. These experiments are consistent with the evolutionary modification of recombinase DNA binding loops to control the on and off rates of recombinases during strand exchange.

### Rad51 is optimized for strand exchange and DNA extension

Substitutions in DNA-binding L2 altered recombination intermediate formation and the extent of genetic recombination. Break-induced replication (BIR) is a type of homology-dependent repair that can be used to complete repair when the second end of DNA lacks homology. This pathway is highly mutagenic but can be beneficial in the recovery of damaged replication forks (Atari *et al*, 2025; Donnianni & Symington, 2013; Lee *et al*., 2024). Our experiments suggested that the *rad51-FD* and *rad51-PD* mutations promote less efficient DNA extension, and we hypothesized that the impact of these mutations may be stronger in assays where repair goes to completion. Therefore, we measured the efficacy of these mutants on BIR outcomes by using a dedicated BIR reporter assay (Anand *et al*, 2014). In this assay system, HO endonuclease induces a double-strand break at a point on chromosome III adjacent to a partial *URA3* gene. The gene sequence can then locate homology at a distal point on the chromosome arm (**Fig. 5A**). This results in strand invasion and the initiation of DNA replication and repair of the broken DNA template. The repair restores a divided URA3 gene, which can be used to select colonies which have used BIR to repair the break. In WT, Rad51 repair happens around 5% of the time (Anand *et al*., 2014). Importantly, this is an endpoint assay that allows us to assay completion of repair.

**Figure 5:**
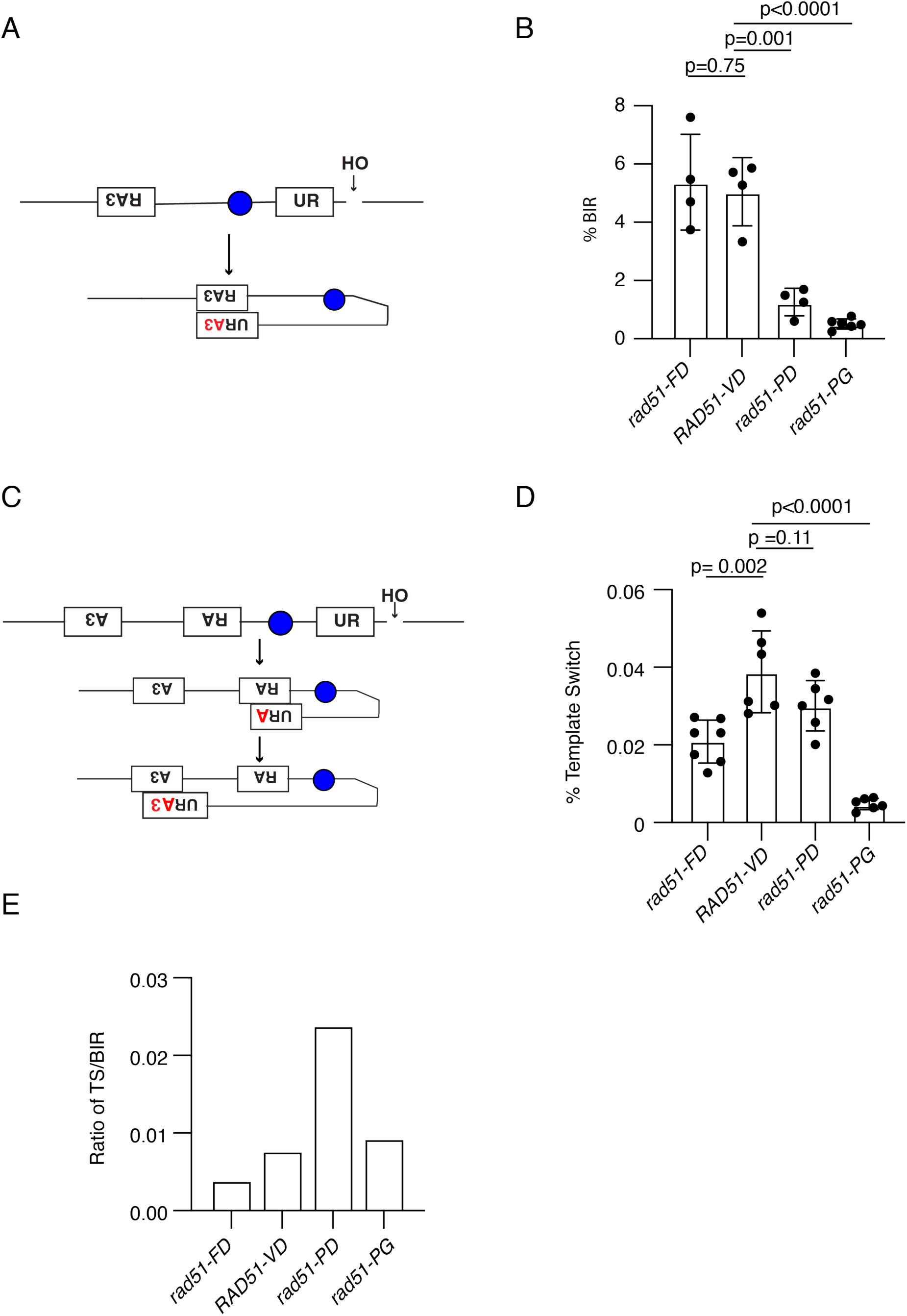
Rad51 is more efficient at BIR but less efficient at template switching. **(A).** Schematic diagram illustrating an assay to monitor the onset of break induced replication (BIR). **(B).** Graph showing the BIR frequency for *rad51-FD, RAD51-VD, rad51-PD,* and *rad51- PG*. The bars represent the mean of the data, and the error bars represent the standard deviation of at least four independent experiments. **(C).** Schematic diagram illustrating an assay to measure the frequency of template switching during BIR. **(D).** Graph showing the Template Switching frequency for *rad51-FD, RAD51-VD, rad51-PD,* and *rad51-PG*. The bars represent the mean of the data, and the error bars represent the standard deviation of at least four independent experiments. **(E).** Graph representing the ratio of BIR to Template switching for *rad51-FD, RAD51-VD, rad51-PD,* and *rad51-PG*.

The *rad51-PD* mutants had a significant 4-fold reduction in BIR efficiency (**Fig 5B**). There was no improved function in *rad51-FD*, and BIR was as efficient as WT (**Fig 5B**). There was an 8–10- fold reduction in survival in the *rad51-PG* strains (**Fig 5B**), consistent with a general reduction in gene conversion observed in the ectopic repair assay, and reduction in D-loop capture efficiency. The defects observed in the *rad51-PD* mutant could be caused by failure to liberate Rad51 from the 3’end of DNA reducing the initiate replication or by accumulation of Rad51 on ssDNA leading to potential toxic recombination intermediates after a migrating bubble has formed (Elango *et al*, 2017; Kramara *et al*., 2018).

A consequence of residual recombinase proteins during DNA extension may be an increased level of Rad51 dependent DNA template switching. To measure this, we used an intrachromosomal template switching assay (**Fig. 5C**) (Anand *et al*., 2014). Like the BIR assay, a DNA break is introduced by expression of the HO endonuclease, and survival is measured as the frequency of *URA3*+ colonies compared to uncut cells. The initial strand invasion is the same as the BIR assay, but the 3’ end of the *URA3* gene is missing. To complete the repair, the initial DNA extension product must dissociate from the template and re-invade a site farther down the chromosome. Completion of this template switch results in cell survival. We measured the ability of Rad51 alleles to survive in a template switching assay. As expected, the *rad51-PG* was reduced by similar amounts as compared to the BIR (**Fig 5D**). In contrast, the *rad51-PD* was not significantly reduced relative to WT (**Fig 5D**). We were surprised to observe that the *rad51-FD* slightly worse at template switching (**Fig 5D**).

Because the -PD mutant is less defective in this assay than in the BIR assay, this implies that it may be more efficient at template switching. A measure of this is the ratio of template switching efficiency/break-induced replication efficiency. In our hands, this ratio is around 0.008 or 0.8%, suggesting that in WT, the template switching reaction occurs in 0.7% of BIR outcomes (**Fig 5E**). The *rad51-PG* mutants were comparable to WT by this metric. In contrast, in the *rad51-PD* mutants, the template switching efficiency was 2.4% (**Fig 5E**), and the *rad51-FD* was around 0.4% (**Fig 5E**). These data suggest that while *rad51-PD* is impaired for BIR, it is more efficient at generating template switching outcomes. This observation is consistent with recombinases that are more difficult to remove from recombination intermediates. In contrast the *rad51-FD* mutant is as efficient at BIR as the WT but is less efficient a template switching. This fits the trend observed for the residues at this position and may be due to the increased rate in D-loop formation observed in the *rad51-FD* mutant, leading to kinetic differences in the repair reaction.

### ATPase and dissociation from dsDNA

Our data suggested that *rad51-PD*, and *rad51-PG* may be more difficult to remove from DNA. We hypothesized that this may be due to a reduction in the ATP hydrolysis rate. Therefore, we purified Rad51, mCherry-Rad51, mCherry-Rad51-FD, mCherry-Rad51-PD, and mCherry-Rad51- PG (**Fig 6A**). For the mcherry-tagged Rad51, the fluorescent protein was in a safe harbor site that allows labelling of Rad51 but does not interfere with activity *in vivo* or *in vitro* (Anand *et al*., 2014; Deveryshetty *et al*, 2025). We compared the ATP hydrolysis rates of unlabeled and labeled Rad51 on ssDNA, and there was no difference in the rate of ATP hydrolysis (**Fig 6B**). We then tested each mutant for their ability to hydrolyze ATP on both ssDNA and dsDNA (**Fig 6B C**). Consistent with our hypothesis, the Rad51-PD and Rad51-PG proteins hydrolyzed ATP ∼2-fold less efficiently than WT (**Fig 6B C**) on both ssDNA and dsDNA. This observation is consistent with altered stability. We also observed a reduction in ATP hydrolysis associated with the Rad51-FD mutant (**Fig 6B C**). This was 75-80% of WT but was not statistically significant.

**Figure 6:**
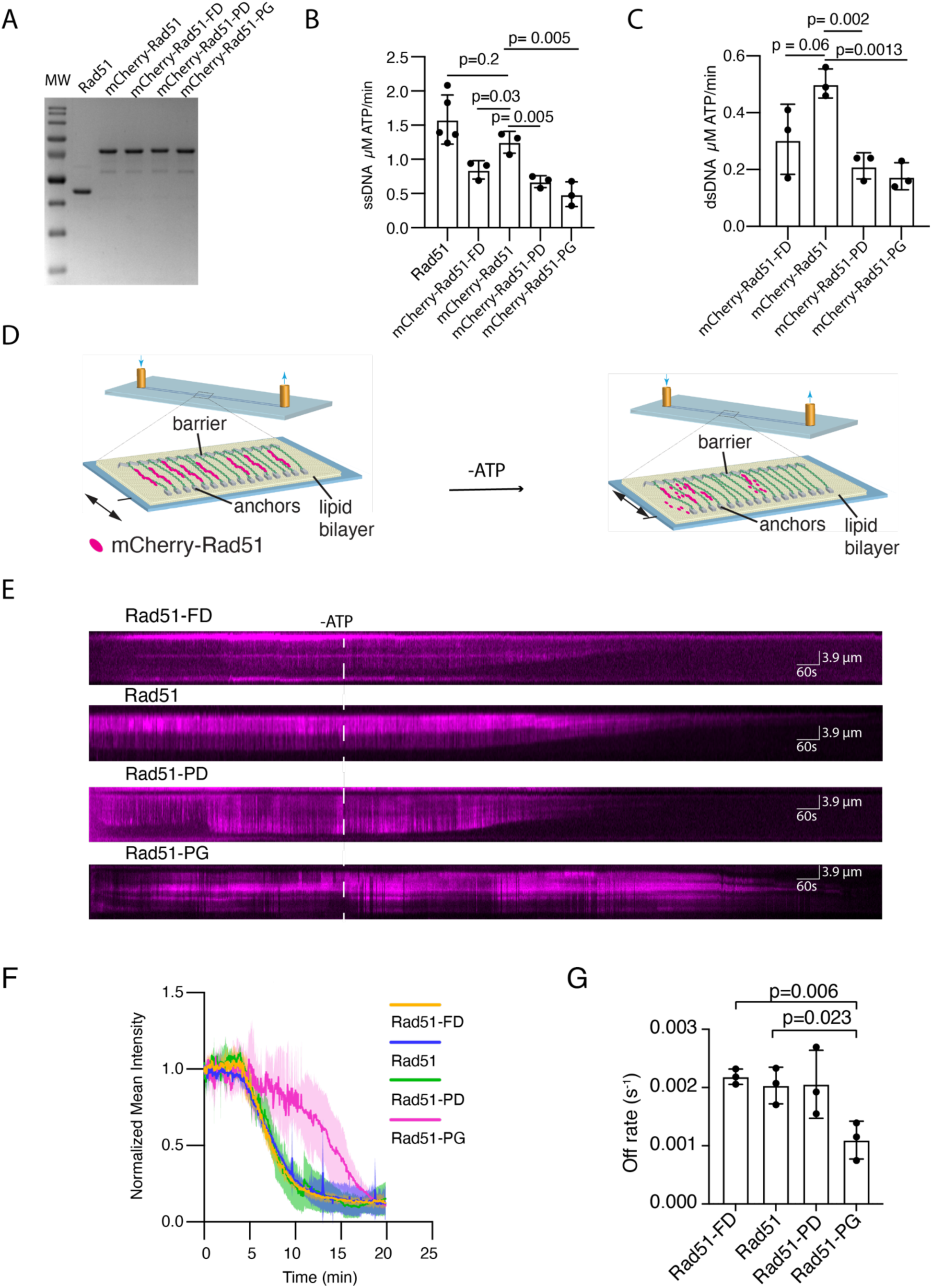
Loop 2 mutants are stable on DNA. **(A).** Representative SDS-PAGE illustrating purification of Rad51, mCherry-Rad51, mCherry- Rad51-FD, mCherry-Rad51-PD, and mCherry-Rad51-PG **(B).** Graph representing the ATP hydrolysis rates for Rad51, mCherry-Rad51, mCherry-Rad51-FD, mCherry-Rad51-PD, and mCherry-Rad51-PG on ssDNA. The bar represents the mean, and the error bars represent the standard deviation of at least three independent experiments. **(C).** Graph representing the ATP hydrolysis rates for mCherry-Rad51, mCherry-Rad51-FD, mCherry-Rad51-PD, and mCherry- Rad51-PG on dsDNA. The bar represents the mean, and the error bars represent the standard deviation of at least three independent experiments. **(D).** Schematic illustrating a DNA-curtain assay to measure the stability of Rad51 on DNA. **(E).** Example kymographs for mCherry-Rad51, mCherry-Rad51-FD, mCherry-Rad51-PD, and mCherry-Rad51-PG dissociation through ATP depletion. **(F).** Graph representing the dissociation of mCherry-Rad51, mCherry-Rad51-FD, mCherry-Rad51-PD, and mCherry-Rad51-PG after depletion of ATP. The line represents the mean of the molecules across three replicates per condition, and the shaded area represents the standard deviation **(G).** A graph representing the dissociation rates for mCherry-Rad51, mCherry-Rad51- FD, mCherry-Rad51-PD, and mCherry-Rad51-PG from dsDNA curtains. The dots represent the mean of individual molecules from at least three independent experiments. The error bars represent the standard deviation.

ATP hydrolysis is a key feature in the disruption of Rad51 filaments bound to ds or ssDNA (Hilario *et al*, 2009; Robertson *et al*, 2009; van Mameren *et al*, 2009). However, the Rad51 protein still has additional affinity for DNA, and hydrolysis is only the first requirement for dissociation. To measure the stability of Rad51 on dsDNA, we used single molecule DNA curtains experiments (**Fig 6D**) (Collins *et al*, 2014; Crickard, 2023). In these experiments mCherry-Rad51 filaments are allowed to form on dsDNA in the presence of ATP. The ATP is then washed out and the dissociation of Rad51 is monitored in real-time. Two effects were observed during filament dissociation: the loss of mCherry-Rad51 signal and the contraction of the dsDNA (**Fig 6E**). The extension and contraction of the dsDNA is a strong indicator correctly formed Rad51 filaments. We measured the rate of dissociation and found that there was no observable difference between Rad51-PD, Rad51-FD and WT (**Fig 6F G**). In contrast, there was a substantial difference in the dissociation rate between WT and Rad51-PG (**Fig 6F G**). This may occur both through increased rates of ATP hydrolysis as well as changes in dsDNA affinity.

## Discussion

In this study, we developed and tested the hypothesis that duplication of the ancestral eukaryotic recombinase gene allowed specialization of Rad51 and Dmc1 within the eukaryotic lineage. This model provides an alternative to the view that only Dmc1 acquired a novel meiosis specific activity following gene duplication. Our results demonstrate that DNA-binding loops inherited from the archaeal eukaryotic ancestor can be modified to regulate recombination outcomes. Substitutions within DNA-binding L2 altered ATP hydrolysis, filament stability, D-loop formation, repair pathway utilization, and crossover frequency. Together, these findings support the idea that duplication of the ancestral recombinase allowed specialization of existing properties, producing a Rad51 optimized for mitotic DNA repair and a Dmc1 adapted to promote genetic exchange during meiosis.

### Rad51 underwent continued specialization following gene duplication

Rad51 has traditionally been regarded as the ancestral eukaryotic recombinase because it performs the conserved function of repairing DNA damage during vegetative growth, a universally conserved HR function. Dmc1 has been viewed as a meiosis-specific innovation, and that duplication of the ancestral gene allowed Dmc1 to specialize for meiosis. Our phylogenetic analyses identified Dmc1-like DNA-binding loop motifs within all archaeal lineages, including members of the Asgard archaea that are thought to be the closest extant relatives of eukaryotes. These observations indicate that amino acid features associated with Dmc1 did not arise de novo following gene duplication in eukaryotes but instead existed within the diversity of archaeal recombinases before the emergence of eukaryotes. Gene duplication in eukaryotes appears to have allowed Dmc1 to develop properties that were already present within an ancestral recombinase.

Consistent with this interpretation, ancestral sequence reconstruction predicted that the last common ancestor of Rad51 and Dmc1 possessed a canonical DNA-binding loop containing the proline-aspartic acid (PD) motif that remains widespread throughout Archaea. The reconstructed protein occupied an intermediate position between modern Rad51 and Dmc1 in overall sequence space, suggesting that it cannot be viewed simply as an ancestral Rad51 or ancestral Dmc1. Instead, the ancestral recombinase likely combined features that were subsequently optimized in distinct directions following duplication (**Fig 7**). This model provides a straightforward evolutionary explanation for why both modern recombinases retain the same core strand exchange chemistry while exhibiting distinct biological functions.

**Figure 7.**
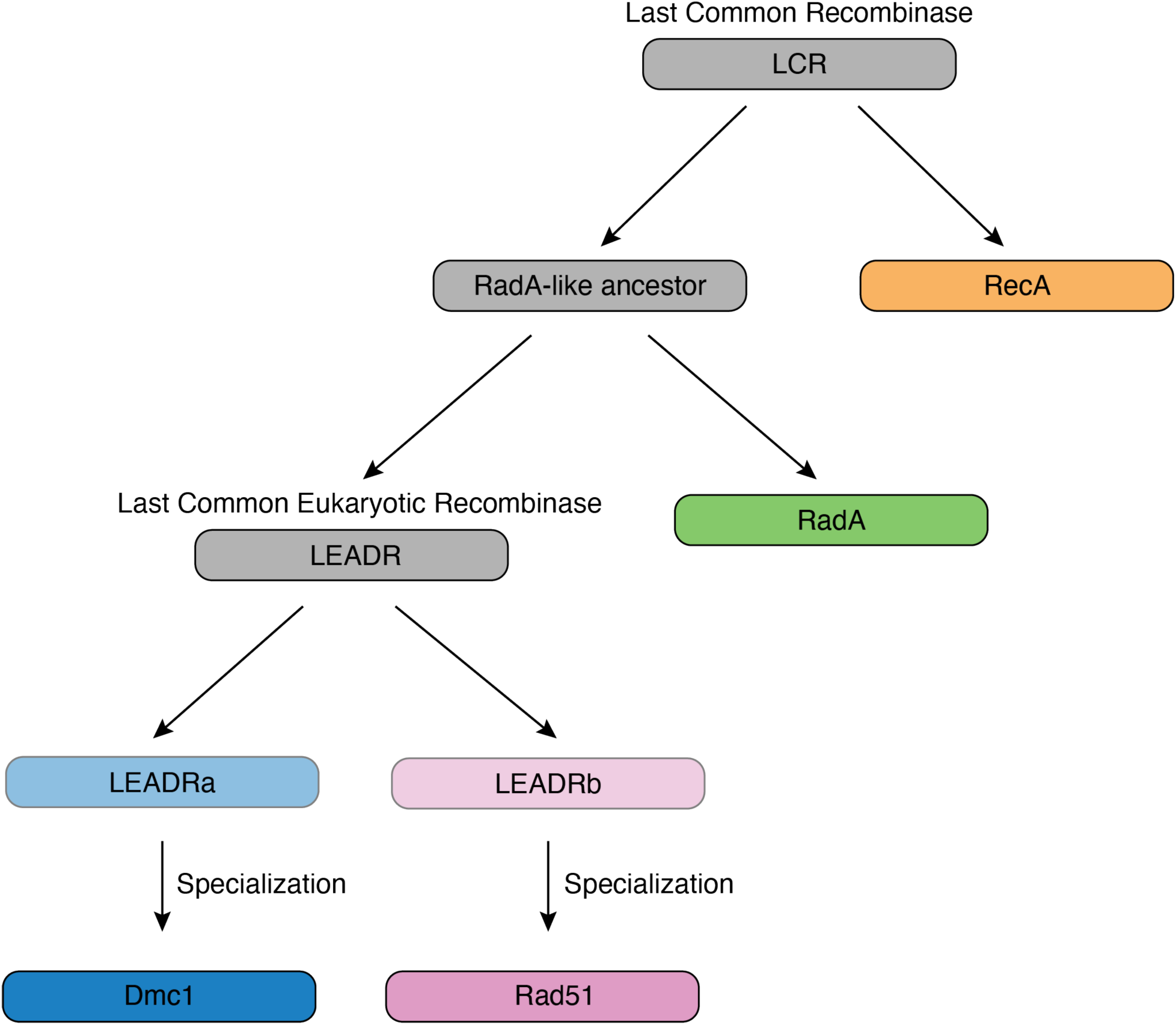
Conceptual model for the evolutionary diversification of RecA-family recombinases. The schematic illustrates the proposed inheritance and diversification of recombinases from a last common recombinase (LCR), giving rise to bacterial RecA and a RadA-like ancestor of archaeal and eukaryotic recombinases. The RadA-like lineage subsequently gave rise to archaeal RadA and the last common eukaryotic recombinase (LEADR). Duplication of LEADR generated two post-duplication ancestral lineages (LEADRa and LEADRb), which subsequently specialized toward Dmc1 and Rad51, respectively. The diagram is intended to represent evolutionary relationships and functional diversification rather than specific ancestral sequence reconstructions.

Following duplication, Rad51 continued to diverge through the acquisition of a valine residue within DNA-binding L2. These changes improved Rad51 for efficient DNA repair, likely enhancing the kinetics of the recombination reaction. Interestingly, although previous biochemical studies suggested that the proline-to-valine substitution in Rad51 slows strand exchange *in vitro*, our *in vivo* experiments demonstrate that it accelerates D-loop formation. One explanation is that purified strand exchange assays do not recapitulate the competitive genomic environment encountered during homology search. Within cells, altered recombinase binding dynamics may facilitate more rapid sampling of homologous sequences, allowing Rad51 to locate productive donor templates more efficiently despite a reduction in intrinsic strand exchange activity.

Our data further suggest that increased ATP hydrolysis alone does not explain the improved repair phenotypes. Although ATP turnover contributes to filament remodeling, the *rad51-FD* mutant exhibited enhanced D-loop formation despite reduced ATPase activity relative to wild type. Likewise, the ancestral *rad51-PD* mutant displayed a modest reduction in ATP hydrolysis yet exhibited defects in DNA extension and break-induced replication that were disproportionate to its effects on D-loop capture. These observations argue that DNA-binding L2 regulates recombinase behavior through a combination of ATP turnover and affinity for DNA. We therefore favor a model in which evolutionary changes within the DNA-binding loops tune the overall dynamics of Rad51 filaments, balancing rapid homology search with efficient filament removal following strand invasion, like that observed for eubacterial RecA (van der Heijden *et al*, 2008).

### Slower strand pairing promotes genetic exchange

An observation from this study was the inverse relationship between D-loop formation and crossover frequency. Variants that promoted rapid D-loop formation consistently produced fewer crossover products, whereas variants that delayed D-loop formation increased crossover formation. This relationship was maintained across all the loop substitutions examined, suggesting that recombinase dynamics can tune the probability of crossover formation. These observations imply that the efficiency of HR is not determined solely by whether strand invasion occurs, but also by how rapidly productive recombination intermediates are formed and subsequently remodeled.

The molecular basis for this relationship remains unresolved. One possibility is that slower strand pairing permits the formation of longer-lived recombination intermediates that are more likely to mature into crossover products. Consistent with this interpretation, the *rad51-PG* mutant produced the strongest defect in D-loop formation while simultaneously exhibiting the greatest increase in crossover frequency and the largest increase in filament stability measured by single-molecule imaging. In contrast, the *rad51-FD* mutant enhanced D-loop formation while virtually eliminating detectable crossover products, demonstrating that efficient homology search and crossover control can be genetically separated. These findings suggest that the timing of early recombination events influences the subsequent choice of repair outcome.

This interpretation is consistent with the activities of anti-recombination factors such as Srs2 and Sgs1, which suppress crossovers by limiting Rad51 filament length and destabilizing recombination intermediates (Ira *et al*., 2003). Although the DNA-binding loop substitutions characterized here are unlikely to regulate the same molecular step as these helicases, both classes of perturbations ultimately alter the lifetime of recombination intermediates and thereby influence pathway choice. Rather than changing the chemistry of homologous pairing, evolutionary modification of recombinase DNA affinity may therefore provide an efficient means of tuning the balance between conservative DNA repair and genetic exchange.

It should be noted that the crossovers observed during mitotic repair are likely distinct from those observed during meiosis, and meiotic crossovers have evolved over millions of years to form with high efficiency. However, crossovers formed during mitosis may represent a pre-cursor state or sub-type of crossovers formed during meiosis and serve as a reasonable proxy for evaluation of genetic exchange for our study.

## Conclusion

Our findings support a revised model for the evolution of eukaryotic recombinases. Rather than Dmc1 alone acquiring a novel function following gene duplication, the ancestral recombinase likely possessed properties intermediate between modern Rad51 and Dmc1. Subsequent specialization of Rad51 produced a more dynamic recombinase that promotes efficient homology search, rapid completion of DNA repair, and faithful recovery of damaged replication forks. In contrast, Dmc1 developed DNA-binding loop properties that stabilize strand exchange intermediates and favor genetic exchange during meiosis.

## Materials and Methods

### Retrieval and Phylogenetic Analysis of Archaeal Recombinase Sequences

Amino acid sequences of putative Asgard recombinases were obtained using BLASTp with *S. cerevisiae* Rad51 (Uniprot P25454), *H. sapiens* Rad51 (Uniprot Q06609), and *S. solfataricus* RadA (Uniprot Q55075) sequences as queries to the Asgard Group database (TaxId: 1935183). The expect value was 0.05 for all searches, and the maximum number of sequences was set to 5,000; for each search, less than 600 hits were returned, all were combined, and duplicate hits were removed. From this set of 653 unique sequences, those shorter than 150 (N = 20) or longer than 500 (N = 5) amino acids were removed, along with 38 sequences that appeared to be partial/incomplete, contained large insertions, or had non-amino acid characters; the final number of sequences used for analysis was 590. Next, we added RecA (*E. coli*) as an outgroup, as well as six RadB reference sequences to aid in the identification of potential RadB homologs in the Asgard group. Sequences were aligned using MUSCLE and manually trimmed to span only the core RecA domain (I41 to G268, *E. coli* numbering). After manual trimming, the alignment was automatically trimmed with BMGE with default settings and a minimum block size of 1.

The trimmed alignment was then used to generate a maximum-likelihood phylogenetic tree to reduce the total sequences in our analysis with Treemmer. For this tree, IQ-tree’s Model Finder selected the LG+R8 substitution model based on the best fit of the Bayesian information criterion. Next, Treemmer was used to reduce the sequences in our dataset while maintaining 95% of the overall tree length. After adding back RadB reference sequences (N = 6) and BLAST hits from Asgard members manually identified as having putative species/strain designations (N = 25), the final number of sequences was 373. The full-length sequences in this dataset were then realigned with muscle, manually trimmed to the core RecA domain using the *E. coli* RecA outgroup as a reference and automatically trimmed with BMGE following the same processes and parameters described above. This dataset’s maximum-likelihood phylogenetic tree was constructed in IQ-Tree with 1000 SH-aLRT (Nguyen *et al*., 2014) and 1000 UFBS (Hoang *et al*., 2017) replicates and the LG + R5 substitution model selected by ModelFinder. We then analyzed and annotated the tree.

To create the RadA-like subtree, the 20 RadB-like hits and 6 RadB reference sequences were first removed from the data set. The remaining 347 sequences were processed following the same steps as described above for sequences that were aligned in MUSCLE, manually trimmed to RecA-core domain, trimmed with BMGE, and used to generate a maximum-likelihood tree with IQtree with E. coli RecA as the outgroup. The substitution model used was LG + R6 as selected by IQ-TREE’s Model Finder. Eighteen groups of putative RadA homologs were assigned based on the criteria that their clade contained four or more sequences and had SH-aLRT/UFBS branch support of >80 and >90, respectively. Of these initial 18 groups, four (groups 2, 3, 14, and 15) lacked clear Walker A, DNA binding Loop 1 and/or DNA binding Loop 2 motifs and were not analyzed further as they are unlikely to represent RadA homologs; similarly, nine individual sequences from the remaining 14 groups also appeared to lack canonical recombinase motifs, or were incomplete, and were removed from our analysis, resulting in 198 Asgard recombinase sequences across 14 groups.

### Collection and analysis of recombinase reference sequences from Archaea, Bacteria, and **Eukarya**

#### Non-Asgard Archaea (RadA)

Sequences for RadA homologs from non-Asgard Archaea were obtained following the same procedure used to recover homologs from the Asgard superphylum. Briefly, *S. solfataricus* RadA, *S. cerevisiae* Rad51, and *H. sapiens* Rad51 were used as queries in BLASTp searches against each archaeal superphyla (TACK Taxid: 1783275; Euryarchaeota (EURY) Taxid: 28890; DPANN Taxid: 1783276). The E-value for each search was 0.05, and the top 500 hits were collected, combined, and duplicate sequences removed. Outlier sequences >500 amino acids long, missing Walker A/B motifs, or containing X characters were removed from each data set. Five outliers were removed for TACK, and the final number of sequences used for all analyses was 758. For EURY, 13 outliers were removed, and the final number of sequences used for all analyses was 782. For DPANN, two outliers were removed, and the final number of sequences used for all analyses was 621. For each superphylum, sequences were aligned with MUSCLE using the default settings in Jalview; logos were generated with Skylign (http://skylign.org).

#### Bacteria (RecA)

Eighty bacterial RecA sequences spanning 40 bacterial phyla (https://lpsn.dsmz.de/text/names-of-phyla) (Oren & Garrity, 2021) were obtained from Uniprot and aligned with MUSCLE using the default settings in Jalview; logos were generated with Skylign (http://skylign.org).

#### Eukarya (Rad51 and DMC1)

The 43 Rad51 and 28 DMC1 sequences used in phylogenetic analyses were obtained from Uniprot and aligned with MUSCLE using the default settings in Jalview; logos were generated with Skylign (http://skylign.org).

### Combined alignment and phylogenetic analysis of bacterial, archaeal, and eukaryotic **recombinases**

A combined dataset (N_total_ = 2511) of Asgard (N = 198), TACK (N = 758), EURY (N = 782), DPANN (N=621), eukaryotic Rad51 (N = 43), eukaryotic Dmc1 (N = 28), and bacterial RecA (N = 81) were aligned with *E. coli* RecA using MAFFT (Katoh *et al*, 2019; Kuraku *et al*, 2013) (https://mafft.cbrc.jp/alignment/server/index.html) with the G-INS-1 progressive method strategy and default settings. The alignment was trimmed manually to the core RecA domain and with BMGE as previously described for all other alignments. IQ-Tree was used to generate a maximum-likelihood phylogenetic tree with 1,000 SH-aLRT and 1,000 UFBS replicates using the LG + R10 substitution model selected by ModelFinder. The tree was visualized and annotated with FigTree and Adobe Illustrator.

### Amino acid pair-frequency analysis

Amino acid frequencies were analyzed at the two multiple-sequence-alignment columns corresponding to *S. cerevisiae* Rad51 residues 331 and 332. Archaeal sequences were grouped into Asgard, TACK, Euryarchaeota (EURY), and DPANN lineages according to the lineage annotations in the sequence headers. For each position, the frequency of an amino acid was calculated relative to the number of sequences containing a valid residue at that position. Gaps and ambiguous residues were excluded from the corresponding denominator. Amino acid pair frequencies were calculated using only sequences containing valid residues at both positions:

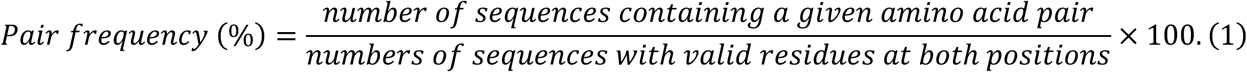

The 15 most frequent amino acid pairs were displayed, with all remaining combinations retained in the denominator. The custom code used for this analysis is provided with the manuscript.

### Ancestral Reconstruction

#### Ancestral sequence reconstruction and sensitivity analysis

The initial sets of eukaryotic Rad51 and Dmc1 sequences and archaeal RadA sequences were obtained from a previous phylogenetic study (Matsuo *et al*., 2025). This data set was distinct from that used to generate the original phylogenetic tree. *S. cerevisiae* Rad51 and *S.s pombe* Rhp51 sequences obtained from EukProt v3 (Richter *et al*, 2022) were added to expand the sampling of the eukaryotic Rad51 clade. Multiple sequence alignments were generated using MUSCLE v5.1 (Edgar, 2022) or MAFFT with the G-INS-i strategy (Kuraku *et al*., 2013). Each alignment was either retained without trimming or trimmed using trimAl (Capella-Gutiérrez *et al*., 2009). Maximum-likelihood phylogenetic trees were inferred using IQ-TREE (Wong *et al*., 2026). The best-fitting amino-acid substitution model was selected separately for each alignment using ModelFinder, and branch support was evaluated using 1,000 ultrafast bootstrap replicates and 1,000 SH-aLRT replicates. Each tree was rooted on the branch separating archaeal RadA from the eukaryotic Rad51 and Dmc1 clades, with the root placed at the midpoint of this branch and named Anc_RadA_Euk. The node representing the last common ancestor of all Rad51 and Dmc1 sequences was named Anc_Euk. The most recent common ancestors of the Rad51, Dmc1, and RadA clades were named Anc_Rad51, Anc_Dmc1, and Anc_RadA, respectively. For each analysis, the corresponding alignment, rooted tree, and best-fitting substitution model were used for ancestral sequence reconstruction in IQ-TREE (Wong *et al*., 2026). ASR results obtained from these original, non-curated datasets were used for the Loop II state-transition analysis and the paired Rad51/Dmc1 evolutionary-trajectory analysis.

#### Robustness analysis of the full-length ancestral sequence

A separate curated analysis was subsequently performed to evaluate the robustness of the full-length Anc_Euk reconstruction and to generate candidate sequences for experimental testing. Four clearly truncated Dmc1 sequences—P016459, P008931, P012347, and P005646—were removed before realignment and phylogenetic reconstruction. Both the t datasets were reported in (Matsuo *et al*., 2025) were analyzed. For the larger dataset, alignments were generated using MUSCLE v5.1 and MAFFT G-INS-i. For the smaller dataset, alignments were generated using MUSCLE v3.8 with default settings through the EMBL–EBI server (Madeira *et al*, 2024) and MAFFT G- INS-i. Each alignment was analyzed in both untrimmed and trimAl-trimmed forms, producing eight ASR pipelines in total. Phylogenetic inference, tree rooting, node naming, and ancestral reconstruction were performed as described above.

Homologous positions were mapped to human RAD51 (P068585 in the source dataset) for residue numbering and positional reference. Because removal of an alignment column does not demonstrate that the corresponding residue was absent from the ancestral sequence, full-length inference was based primarily on the four untrimmed alignments. For each untrimmed alignment, columns with *P*(*present*) ≥ 0.90, as inferred using a sitewise presence/absence model, were retained in the exploratory full-length candidate.

The major uncertainties involved the acidic, low-complexity 33-aa N-terminal region, the retention of the position corresponding to human RAD51 H275, and several core positions with closely competing ancestral states. The recommended primary sequence retained the coherent 33-aa N-terminal block reconstructed from the curated larger-dataset MUSCLE v5.1 alignment. At each remaining human-RAD51-anchored coordinate, posterior probabilities for the 20 amino acids were averaged with equal weight across all curated ASR pipelines in which that coordinate was represented, and the amino acid with the highest mean posterior probability was selected. This procedure was treated as a cross-pipeline sensitivity consensus rather than formal Bayesian model averaging because the pipelines were correlated. Alternative sequences represented uncertainty in the N-terminal alignment and at 13 close-tie core positions. The latter were defined as positions with mean top-state support below 0.8 and a difference of less than 0.15 between the top- and second-ranked amino acid states.

#### Loop 2 transition analysis

The two alignment positions corresponding to the focal residues in DNA-binding loop 2 were identified separately in each non-curated alignment. Observed Rad51 and Dmc1 motifs were extracted from tip sequences. For reconstructed internal nodes, the maximum-posterior amino-acid state at each of the two positions was obtained from the IQ-TREE ancestral-state output, and the two states were concatenated to define the reconstructed Loop 2 motif. Motif confidence was conservatively defined as the lower of the two site-specific posterior probabilities. Loop 2 motifs were then mapped onto the rooted Rad51 and Dmc1 subtrees. A transition was assigned whenever the maximum-posterior motif differed between a parent and its direct child node. Detailed transition trees retained all observed tips and reconstructed internal nodes, whereas summary trees contracted contiguous parent-child regions with the same motif so that every remaining edge represented an inferred motif transition. This analysis represents maximum-posterior ancestral-state mapping rather than stochastic character mapping. Unless otherwise indicated, the Loop 2 transition figures were generated from full, non-curated larger dataset using MUSCLE v5.1 and analyzed without alignment trimming. The custom code used for this analysis is provided with the manuscript.

#### Paired Rad51/Dmc1 evolutionary-trajectory analysis

Family consensus sequences and diagnostic sites were defined from the observed Rad51 and Dmc1 tip sequences. At positions with amino-acid occupancy of at least 0.5, the most frequent residue among occupied sequences was assigned as the family consensus state; positions with lower occupancy were treated as missing. A position was classified as Rad51/Dmc1-diagnostic when amino-acid occupancy was at least 0.7 in both families, the modal residue accounted for at least 0.80 of occupied sequences in each family, and the modal residues differed between the two families. Gaps and ambiguous residues were excluded from occupancy-adjusted frequency denominators. Species were included when unique Rad51 and Dmc1 tips could be identified. For each selected sequence, the directed path from Anc_Euk to the corresponding tip was extracted from the rooted tree, including all intervening reconstructed nodes. At each node, the fractions of comparable diagnostic sites matching the Rad51- and Dmc1-type states were calculated, together with pairwise sequence identities to the corresponding extant protein and to the Rad51 and Dmc1 family consensus sequences. Pairwise identities were calculated over alignment positions containing a standard amino acid in both sequences; positions containing a gap or non-standard residue in either sequence were excluded. Unless otherwise indicated, the paired evolutionary-trajectory analysis figures were performed using full, non-curated larger dataset aligned with MUSCLE v5.1 without trimming. Custom code is provided with the manuscript.

#### Yeast strains construction

All yeast strains used in this study can be found in **Supporting Table 2**. Strains for the ectopic recombination assay were TGI354 and were generated by gene replacement with plasmids listed in **Supporting Table 3**. Strains for the *in vivo* D-loop capture (DLC) and extension (DLE) assay were generated from the WDHY5511 parent by gene replacement with plasmids and listed in **Supporting Table 3**. Parent strains for BIR and template switching are from yRA53 (Template Switch) and yRA213 [BIR](Anand *et al*., 2014). In the template switching strain the KANMX cassette was replaced with HPHMX before Rad51 mutations were made by gene replacement with plasmids listed in **Supporting Table 3**. All point mutations were confirmed by sequencing the Rad51 Gene.

#### Ectopic recombination assay

The wildtype strain used for this study was described in (Ira *et al*., 2003). The genotypes for modifying these strains are listed in supporting table 1. A single colony was picked and grown in YP + 3% glycerol + 2% lactate medium to log phase. The culture was diluted and plated on YP + 2% dextrose and YP + 2% galactose plates, respectively. Plates were incubated at 30 °C for 2 to 3 days, and the number of colonies was counted. The viability rate was calculated by dividing the colony number on the YP + 2% galactose plate by that on the YP + 2% dextrose plate.

#### Crossover Analysis

A single colony was picked and grown in YP + 3% glycerol + 2% lactate medium to log phase. The culture was diluted and plated on YP + 2% galactose plates. Plates were incubated at 30 °C for 3 days. Single colonies were picked and resuspended in a spheroplast buffer (400 mM sorbitol, 400 mM KCl, 40 mM Na₃PO₄, 0.5 mM MgCl₂, and 17.5 µg/mL zymolyase). After incubation at 37°C for 30 min, zymolyase was heat-inactivated at 95° for 10 minutes. Spheroplasts were diluted 6.5-fold with DiH_2_O and used as templates for PCR (Primers: P5 5’- CAAGACTGTCAAGGAGGGTATTC-3’, P6 5’-TCAGGGATACCAGCATACTC-3’, P7 5’- GATGCCCTTGTTTTGTTTACTG-3’). PCR products were analyzed by 1% agarose gel electrophoresis.

#### BIR and Template Switching Assays

Individual isolated colonies were grown in 5 mL YP + 3% glycerol + 2% lactate for 16 hours. Cells were then diluted to an appropriate dilution and plated on either YNB (SC) + 2% dextrose or YNB (-URA) + 2% galactose. After 3 days, the number of colonies were counted, and Colony Forming Units (CFU) were calculated for each plate. The CFU from the -URA plate was then divided by the SC plate, and the frequency of BIR or Template Switching was determined.

#### DLC assay

DLC assay was performed as described (Piazza *et al*., 2019; Reitz *et al*., 2022). In brief, yeast cells were grown in a 5 mL YP + 2% dextrose + 4% adenine sulfate medium overnight. The second day, the culture was diluted by 10-fold in 5 mL YP + 3% glycerol + 2% lactate + 4% adenine sulfate and grown for around 8 hours. Then the culture was inoculated into 100 mL of YP + 3% glycerol + 2% lactate + 4% adenine sulfate medium with an initial OD_600_ ≈ 0.006 and grown for 16 hours. A 5x psoralen stock solution (0.5 mg/mL trioxsalen in 200-proof ethanol) was made in a 50-mL aluminum foil-covered falcon tube and dissolved on a shaker at room temperature overnight with gentle rocking. Next day, the culture should have an OD_600_ at 0.3-0.8. 7.5 OD_600_ of cells was collected as time 0 control, centrifuged at 2,246 x g, 4 °C for 5 minutes.

The cell pellets were resuspended in 1x psoralen buffer. The 1x psoralen buffer was prepared by diluting 5x psoralen in TE1 buffer (50 mM Tris-Cl [pH 8.0], 50 mM EDTA) before collecting cells. The resuspended cells were plated in a 60 mm x 15 mm petri dish, put 2-3 cm below a UV light source with the lip removed atop a pre-chilled metal block. The cell samples were exposed under the UV light for 10 minutes with gentle shaking to crosslink DNA. The cells were transferred to a 15-mL Falcon tube. The petri dish was rinsed with TE1 buffer, and the TE1 buffer was poured over the cells. The cells were then centrifuged at 2,246 x g, 4 °C for 5 minutes again. The supernatant was properly disposed of, and the pellets were stored at -20 °C. Galactose was added to the culture to a final concentration of 2% to induce DSBs. Cells were collected at the designed time points as described above. For lysis, the cell pellets were thawed on ice, then resuspended in spheroplasting buffer (0.4 M sorbitol, 0.4 M KCl, 40 mM sodium phosphate buffer [pH 7.2], 0.5 mM MgCl_2_) and transferred to a 1.7 mL microfuge tube.

The cells were spheroplasted in zymolyase solution (2% glucose, 50 mM Tris-Cl [pH 7.5], 5 mg/mL zymolyase 100T) at 30 °C for 20 minutes. The cells were washed with spheroplasting buffer three times at 2,500 x g and restriction enzyme buffer (RE buffer: 50 mM potassium acetate, 20 mM Tris-acetate, 10 mM magnesium acetate, 1 mg/mL BSA) at 16,000 x g three times. The pellets were resuspended in 1.4x RE buffer, either alone or with a hybridization oligo to restore the *EcoR*I restriction sites. The DNA was solubilized by incubating the cells with 0.1% SDS at 65 °C for 13 minutes. 1% Triton X-100 quenched the SDS. The DNA was digested by 20 U *Eco*RI at 37 °C for 1 hour. The restriction enzyme was deactivated by incubating the DNA with 1.5% SDS at 55 °C for 10 minutes. The cells were returned to ice, and SDS was quenched by the addition of 6% Triton X-100. Ligation buffer (50 mM Tris-HCl [pH 8.0], 10 mM MgCl_2_, 10 mM DTT, 2.5 μg/mL BSA, 1 mM ATP, pH 8.0, 8 U T4 DNA ligase) was added to perform the ligation reaction at 16 °C for 1 hour and 30 minutes. 25 μg/mL protease K was added to digest the enzymes at 65 °C for 30 minutes. DNA was extracted by adding phenol:chloroform:isoamyl alcohol and vortexing. The upper water phase was moved and incubated with a tenth volume of sodium acetate and a volume of isopropanol at room temperature for 30 minutes and centrifuged at 21,130 x g, 4 °C for 10 minutes to get DNA precipitation. The DNA pellets were washed with 70% ethanol, dried at 37 °C and then dissolved by incubating with 1xTE buffer (10 mM Tris-Cl [pH 8.0], 1 mM EDTA) at 37 °C for 1 hour. The DNA was used as a qPCR template with the primers listed in (**Supporting Table 4**). DLC chimera content was calculated by [DLC amplification efficiency]^[- Cp_(DLC)_], and the intramolecular ligation product content was calculated by [intramolecular ligation amplification efficiency]^[-Cp_(ligation)_]. The final DLC signal was calculated as DLC chimera content divided by intramolecular ligation product content.

#### DLE assay

DLE assay was performed as described before(Piazza *et al*., 2019; Reitz *et al*., 2022). The procedures were similar with DLC assay except for (1) 2.5 OD_600_ of cells were collect at each time point; (2) DNA crosslinking was omitted and cell pellets were washed by TE1 buffer twice at 2,246 x g; (3) hybridization oligos were different and listed in Supporting Table 3; (4) DNA was solubilized by 1% SDS at 65 °C for 15 minutes; (4) DNA digestion was performed by adding 20 U *Hin*dIII; (5) qPCR was performed using some different oligos listed in Supporting Table 3.

#### Rad51 Purification

WT Rad51 was purified as previously described (Crickard *et al*, 2020). mCherry-Rad51 and all mutants were overexpressed in *E. coli* BL21 (DE3) Rosetta2. For expression, 1.5 L of cells were grown at 37 °C to an OD_600_ of 0.6–0.8 before being induced by 0.5 mM IPTG for 16 hr at 18 °C. Then, the cells were harvested and washed with Lysis Buffer (25 mM Tris-Cl [pH 7.5], 200 mM NaCl, 10% glycerol, 0.05% Igepal-680, and 4.9 mM 2-mercaptoethanol). The cells were lysed by freezing as pellets at -80 °C. The frozen pellet was thawed and resuspended in Lysis Buffer supplemented with protease inhibitor cocktail (Roche Cat. No. 05892953001) and 2 mM PMSF. The resuspended cells were lysed by sonication on ice for ten 15 s/min duty cycles at 65% amplitude. After sonication, the lysate was centrifuged at 17,096 x g for 45 min, and the supernatant was filtered through a 0.22 µM PVDF filter. Purification was conducted using the fast liquid protein (FLP) chromatography ÄKTA go system (Cytiva) and controlled by Unicorn 7.11, the software supplied by the manufacturer. A low salt Buffer A (25 mM Tris-Cl [pH 7.5], 10% glycerol, 0.05% Igepal-680, and 4.9 mM 2-mercaptoethanol (BME)) and a high salt Buffer B (25 mM Tris-Cl [pH 7.5], 1 M NaCl, 10% glycerol, 0.05% Igepal-680, and 4.9 mM BME) were used in this procedure. Two chromatographies were done. In the first chromatography, a heparin column (HiPrep™ Heparin FF 16/10, Cat. No. 28936549) was equilibrated with 2 column volume (CV) of an 80% Buffer A, 20% Buffer B mixture and the cell extract was applied to it at 2 mL/min. After sample application, the column was washed with 2 CV of 80% Buffer A, 20% Buffer B. Proteins were eluted from the column by gradient elution using 10 CV of 20%–100% Buffer B in Buffer A. 3 mL elution fractions were collected and fractions containing the protein of interest were determined via A280 absorption and SDS-PAGE with Coomassie Blue staining. These peak fractions were pooled together and dialyzed using SnakeSkin™ (Thermo Scientific Cat. No. 68100) in 20% Lysis Buffer for 16 hours at 4 °C (in preparation for another chromatography. In the second chromatography, a Q Sepharose column (HiScreen™ Q HP, Cat. No. 28950511) was equilibrated with 2 column volume (CV) of an 80% Buffer A, 20% Buffer B mixture and the cell extract was applied to it at 1 mL/min. The column was washed and eluted in the same manner as the previous chromatography. As previously described, 3 mL peak fractions were collected and pooled. Proteins were concentrated to 6 mg/mL using a 10 kDa cut-off column, Vivaspin (Cytiva Cat. No. 28932296). Concentrated proteins were stored at -80 °C.

#### ATP hydrolysis assay

A commercially available ADP-GLO kit was used to measure ATP hydrolysis rates. The ATP hydrolysis reaction was performed in HR buffer (20 mM Tris-Cl [pH 7.5], 50 mM NaCl, 10 mM MgOAc_2_, 200 ng/µL BSA, 5 mM BME, 100 µM ATP, and 10% Glycerol) and contained 0.1 mg/mL of Phi X circular ssDNA or Phi X RFII and 1 µM recombinase. Reactions were conducted at 30 °C for 20, 40, and 60 minutes. Chemiluminescence was measured using an Azure 400 Fluorescent Imager.

#### Flow cell construction

Metallic chrome patterns were deposited on quartz microscope slides with predrilled holes for microfluidic line attachment by electron beam lithography to generate flow cells. After metal deposition, a channel was created by placing a small piece of paper between the drill holes and covering the two-sided tape. The paper was excised to make the flow chamber, and a glass coverslip was fixed to the tape. The chamber was sealed by heating to 165 °C in a vacuum oven at 25 mmHg for 70 minutes. Flow cells were then completed by hot-gluing IDEX nano ports over the drill holes on the opposite side of the microscope slide from the coverslip.

#### Single-molecule experiments

All single molecule experiments were conducted on a custom-built prism-based total internal reflection microscope (Nikon) equipped with a 488-nm laser (Coherent Sapphire, 100 mW), a 561- nm laser (Coherent Sapphire, 100 mW), a 640-nm laser (Coherent Obis, 100 mW), and two Andor iXon EMCCD cameras. DNA substrates for DNA curtains experiments were made by attaching a biotinylated oligo to one end of the 50 kb Lambda phage genome and an oligo with a digoxigenin moiety on the other. For the purposes of this experiment, only the biotinylated end was used to single tether the DNA on the chrome barriers using streptavidin. Flow cells were attached to a microfluidic system, and sample delivery was controlled using a syringe pump (KD Scientific). Three-color imaging was achieved by two XION 512 × 512 back-thinned Andor EM-CCD cameras.

The ATP-depletion induced dissociation rates of mCherry tagged Rad51 variants were measured by initiating data collection, followed by the injection of 3.6 µM Rad51 in Curtains HR buffer (30 mM Tris-OAc [pH 7.5], 10 mM Mg(OAc)_2_, 50 mM NaCl, 0.5 mM DTT, 2 mM ATP, 0.2 mg/mL BSA) onto single-tethered DNA curtains at a flow rate of 1 mL/min for 12 seconds to allow proteins to bind to DNA. This was followed by a 5-minute incubation step with no flow and a 5- minute wash step with Curtains HR buffer at 0.2 mL/min. Then, the flow was switched from Curtains HR buffer to ATP Depletion HR buffer (Curtains HR Buffer without ATP) and data collection continued for another 20 minutes. Images were collected with a 200 msec integration time at 2 sec intervals. The photobleaching rate of the laser on the mCherry fluorophore was controlled for by using the same experiment but with 600 nM Rad51 and by continuing with Curtains HR Buffer for an additional 12 minutes instead of switching to ATP Depletion HR Buffer. Images were collected with a 200 msec integration time at 200 msec intervals for the final 12 minutes for the bleaching experiment.

For the data analysis, kymographs were drawn on 1 x 50 pixel regions of interest (ROIs) on the long axis of individual dsDNA molecules on the TIFF stacks of the experiment. Intensity data was extracted out of these kymographs and decay was analyzed using homemade MATLAB scripts. These scripts took in the kymographs as inputs as well as hand drawn ROIs to determine the background to normalize to and to determine where the Rad51 filaments were to determine total filament intensity for each frame. Decay was calculated starting with the first frame after flow switches to the ATP Depletion HR Buffer with the intensity of the filament normalized to the mean of the first three frames after the wash step had ended. The decay curves of the individual kymographs emerging from the same experiment were pooled together to generate a mean curve for analysis. The exponential function *A= A_max_e^-kt^* was used to fit the fluorescent signal decay in both experiments. The dissociation rate was determined by calculating the difference between the *k* values obtained from a bleaching experiment and dissociation experiment: *k_off_ =k_dissociation_ - k_bleaching_*.

## Data availability

All primary data will be uploaded to a Mendeley (https://data.mendeley.com/preview/k2w46jwtc8?a=d8525ab1-1e10-40ef-8980-2fe28bae6e59) repository upon acceptance of this manuscript

## Code availability

Code and workflow will be available on-line upon acceptance of this manuscript

## Author contributions

BF and JH performed bioinformatic analysis and JH performed Ancestral sequence reconstruction, PH performed biochemical experiments, analyzed data, and helped write the manuscript. JH performed D-loop capture and extension experiments and analyzed data. IK performed and analyzed all single molecule experiments. JBC provided reagents, performed experiments, guided experiments, and wrote the manuscript with input from BF, PH, IK and JH. NIGMS R35142457 supported this work to JBC.

## Declaration of AI

No generative AI was used in the development of this manuscript. AI was used to refine code and during the editing of the manuscript. AlphaFold was also used in generation of predicted protein structures.

## Supporting information

Supplemental_FiguresandTables

## Acknowledgments

We would like to acknowledge Thorsten Allers, John Logsdon Jr., and Jim Haber for their comments on the study, Eric Alani for carefully reading the manuscript, and members of the Crickard laboratory for reading it. We also acknowledge that original strains used in these experiments were a kind gift from Wolf Heyer and Jim Haber, and the plasmid for expressing and purifying labeled Rad51 was a gift from Edwin Antony. We would also like to thank members of the Cornell R3 groups for their comments during the development of this project.

