## Supplemental_FiguresandTables for "Evolutionary specialization of Rad51 and Dmc1 tune recombination outcomes"

### Extended View:

**Extended View Tables S1, S2, S3,S4**
S1,S2, and S3 tables are related to methods
S4 is related to Figure 2 and S7

**Extended View Figures EV1-10**
EV1 and EV2 are related to Figures 1 and 2
EV3  and EV4 is related to Figure 1
EV5-10 is related to Figure 2

**Supporting Table 1**
**Rad51/Dmc1 Diagnostic sites**

| Site* | Rad51_state | Rad51_occupancy | Rad51_frequency | Dmc1_state | Dmc1_occupancy | Dmc1_frequency | Site in *H. sapiens* RAD51 | Site in *H. sapiens* DMC1 | LEADR_state |
|---|---|---|---|---|---|---|---|---|---|
| 1169 | N | 0.986 | 1 | T | 0.817 | 0.968 | 196 | 195 | N[R] |
| 1215 | Y | 0.873 | 1 | F | 0.983 | 0.968 | 228 | 229 | F[D] |
| 1217 | T | 0.901 | 1 | V | 0.817 | 0.968 | 230 | 231 | T[R] |
| 1249 | D | 0.901 | 1 | E | 0.850 | 0.968 | 257 | 259 | E[D] |
| 1254 | G | 0.944 | 1 | N | 0.984 | 0.984 | 260 | 261 | N[D] |
| 1268 | V | 0.859 | 1 | P | 0.983 | 0.968 | 273 | 274 | P[D] |

***: indicate the column number in MSA file.**
**[R]: indicate the state is Rad51-like.**
**[D]: indicate the state is Dmc1-like.**

**Supporting Table 2**
**Strains used in the study**

| Strains | Genotype | Source or reference |
|---|---|---|
| WDHY5511 | *ura3::A-HOcs, lys2::A, trp1::GAL-HO-hphMX, his3D200, can1-100, leu2–3, 112, ade2–1, RAD5* | Piazza et al 2020 |
| JBC423 (WDHY5511) | *RAD51::rad51-V331F-KANMX* | This study |
| JBC281 (WDHY5511) | *RAD51::rad51-V331P-KANMX* | This study |
| JBC367 (WDHY5511) | *RAD51::rad51-V331P, D332G-KANMX* | This study |
| TGI354 | *TGI 354 JKM 146 (arg5,6::MATa-HPH)* | Ira et al 2003 |
| JBC425 (TGI354) | *RAD51::rad51-V331F-KANMX* | This study |
| JBC357 (TGI354) | *RAD51::rad51-V331P-KANMX* | This study |
| JBC359 (TGI354) | *RAD51::rad51-V331P, D332G-KANMX* | This study |
| JBC361 (TGI354) | *RAD51::KANMX* | This study |
| yRA53 | *MATa::Del HOcs::hisG ura3D851 trp1DEL63 leu2DEL::KAN hmlDEL::hisG HMR::ADE3 ade3::GAL::HO can1DEL::UR::HOcs::NAT, RA-LEU2, A3::TRP1 KAN::HPHMX* | Anand et al 2014 |

| | | |
|---|---|---|
| JBC677 (yRA53) | *RAD51::rad51-V331F-KANMX* | This study |
| JBC633 (yRA53) | *RAD51::rad51-V331P-KANMX* | This study |
| JBC603 (yRA53) | *RAD51::rad51-V331P, D332G-KANMX* | This study |
| yRA213 | *MATa::DEL HOcs::hisG ura3DB51 trpDEL.63 leu2DEL::Kan hmlDEL::hisG HMR::ADE3 ade3::GAL::HO can1DEL::URA::HOcs::HPH, A3::TRP1* | Anand et al 2014 |
| JBC668 (yRA213) | *RAD51::rad51-V331F-KANMX* | This study |
| JBC638 (yRA213) | *RAD51::rad51-V331P-KANMX* | This study |
| JBC621 (yRA213) | *RAD51::rad51-V331P, D332G-KANMX* | This study |

**Extended View Table 3**
**Plasmid in the study**

| Backbone | Construction | Source |
|---|---|---|
| pUC19 | *RAD51-KANMX* | This study |
| pUC19 | *rad51-V331F-KANMX* | This study |
| pUC19 | *rad51-V331P-KANMX* | This study |
| pUC19 | *rad51-V331P, D332G-KANMX* | This study |
| | mCherry-RAD51 | This study |
| | mCherry-rad51V331F | This study |
| | mCherry-rad51V331P | This study |
| | mCherry-rad51V331P, D332G | This study |

**Extended View Table 4**
**Oligos in the study**

| Name | Sequence | Purpose |
|---|---|---|
| 90-mer DNA | Atto647N-GATGTTCTGCTGGATATGCACTTTTCCGGGCTGACGTACACCGTGCTCAGCCTGTTTTTCAGCGATCCGGATATGCATCCGCTGGATTTC | Single Molecule imaging |
| olWDH1760 | CAGCGGGCTTGCAGAAGTTG | To amplify genomic DNA at *ARG4* |
| olWDH1761 | GGCCAATTAGTTCACCAAGACG | To amplify genomic DNA at *ARG4* |

| | | |
|---|---|---|
| olWDH1766 | GTTTCAGCTTTCCGCAACAG | To quantify DSB induction |
| olWDH1767 | GGCGAGGTATTGGATAGTTCC | To quantify DSB induction |
| olWDH2009 | CACCACTTTGCCATTCAACAC | To amplify at the downstream site of elongation |
| olWDH2010 | TGCTCGGAGATTACCGAATC | To amplify at upstream site of the DSB, used with olWDH2009 to quantify DLE signal |
| olWDH2011 | TGCGAGGTTTTCTTGGTCAG | Used with olWDH2009 to quantify *Hin*dIII recognition site restoration |
| olWDH2012 | CGAGGCATATTTATGGTGAAGG | Used with olWDH2010 to measures *Hin*dIII recognition site restoration |
| olWDH2052 | ATGTGCCTTCCTACCGCTC | To quantify intramolecular ligation efficiency of *Hin*dIII-derived fragments |
| olWDH2053 | TCAAGCGTGGTTACATTCCTTAC | To quantify intramolecular ligation efficiency of *Hin*dIII-derived fragments |
| olWDH2007 | TCTGCTCGGAGATTACCGAATCAAAAAAATTTCAAAGAAACCGGAATCAAAAAAAAGAACAAAAAAAAAAAAGATGAATTGAAAAGCTTTATGGACCGAC | To restore *Hin*dIII site |
| olWDH2046 | AATCTTTGTGAAGCTTCGCAAGTATTCATTTTAGACCCATGGTGGAACCCTAGTGTTGAATGGCAAAGTGGTGATAGAGTTCATAGAATTGGTCAGTAT | To restore *Hin*dIII site |
| olWDH1762 | ACTTCGAATTTCGGCACTTC | To quantify intramolecular ligation efficiency of *Eco*RI-derived fragments |
| olWDH1763 | CGATGAAACGTTAAGTGACCAC | To quantify intramolecular ligation efficiency of *Eco*RI-derived fragments |

| | | |
|---|---|---|
| olWDH1764 | AGAGCGGTCAGTAGCAATCC | To amplify at the upstream of DSB |
| olWDH1765 | CACACGCGAAAAACCGCC | To amplify at the upstream of the donor DNA, used with olWDH1764 to quantify DLC signal |
| olWDH2019 | CTTTAACCGGACGCTCGA | To quantify psoralen crosslinking efficiency |
| olWDH2020 | TTGAGTTTATTGCTGCCGTC | To quantify psoralen crosslinking efficiency |
| olWDH1768 | AGGAGCACAGACTTAGATTGG | Used with olWDH1764 to measure *Eco*RI recognition site restoration |
| olWDH1770 | CGAAATCATCTTCGGTTAAATCCAAAACGGCAGAAGCCTGAATGAAACATATGAACCAATTGGAGGACGTCAATGAATTCTGGGGATCCATTGCATTTTT | To restore *Eco*RI site |

### Extended View Figure 1

A

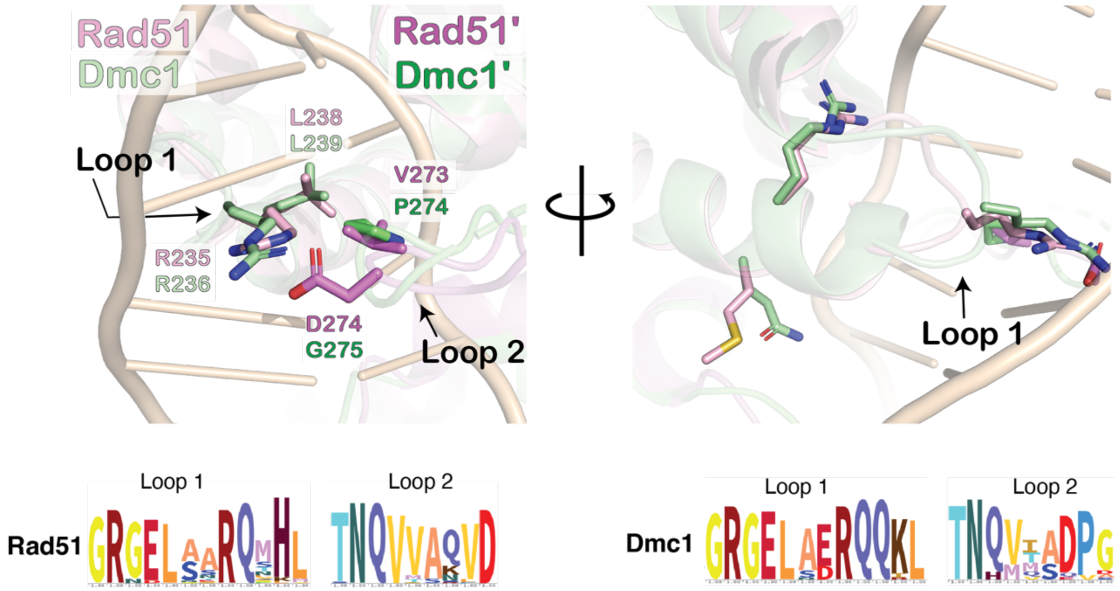


**EVFig 1: Differences in DNA binding loop interactions between Rad51 and Dmc1**
**(A).** Superposition of hRAD51 (PDB:5HC1; protomer 1 in light pink, protomer 2 in violet) and hDMC1 (PDB: 7C98; protomer 1 in palegreen, protomer 2 in lime) structures highlighting the differences in amino acids proposed to regulate mismatches in DNA (also indicated by dashed boxes in amino acid logos) **(Top right)**. Amino acid sequence logos for Rad51 (top, N = 43) and Dmc1 (bottom, N = 28) DNA binding Loop 1 and Loop 2 regions. **(Bottom).**

### Extended View Figure 2

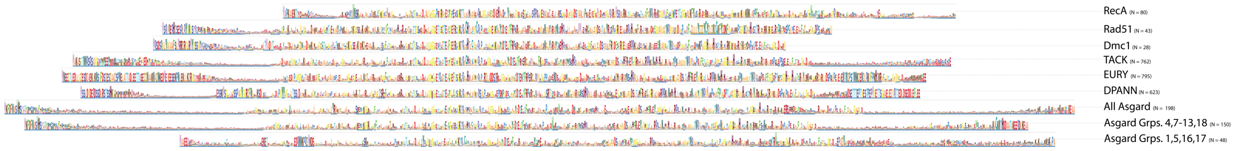


**EVFig 2: Full-length sequence logos for recombinase groups**
Full-length logos used in **Figure 1**: RecA (N = 81), Rad51 (N = 43), Dmc1 (N = 28), TACK (N = 758), EURY (N = 782), DPANN (N = 621), and Asgard (N = 198) for complete dataset.

### Extended View Figure 3

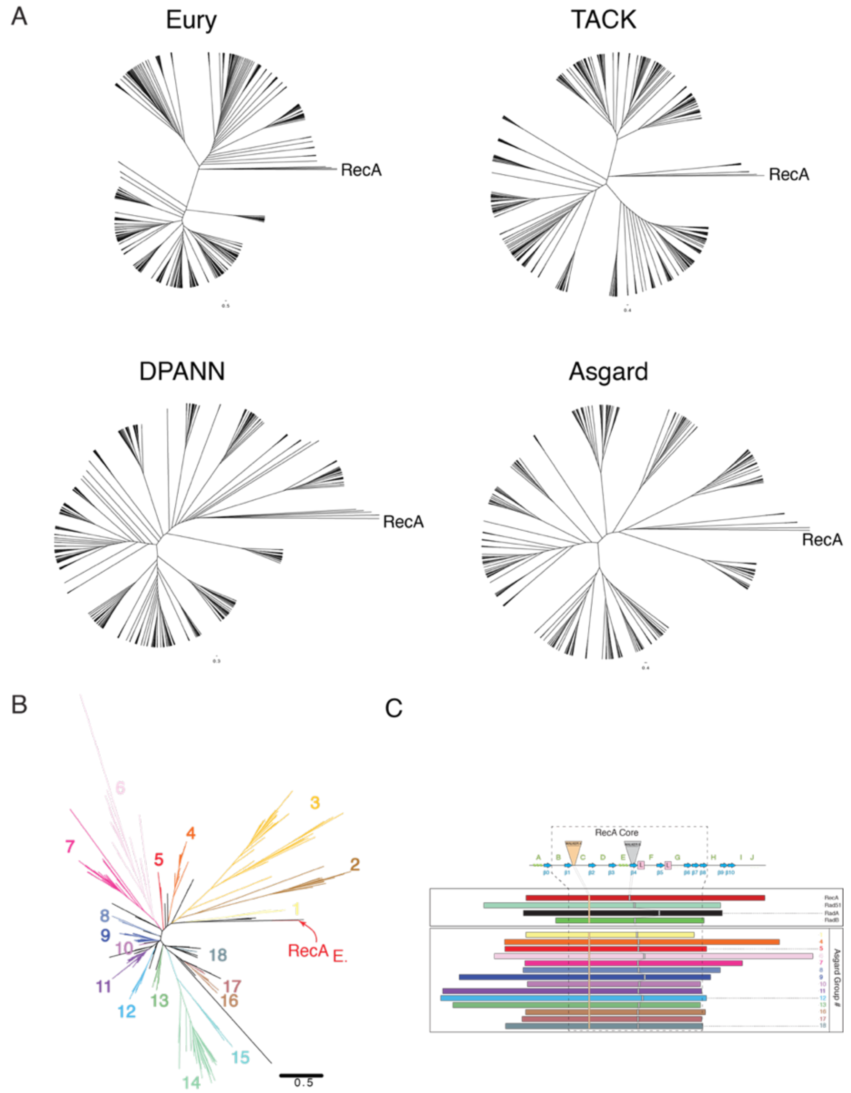


**EVFig 3: Phylogentics of the Archaea Superphylum**

**(A).** Cladogram view of maximum likelihood phylogenetic tree built from recombinase sequences for the EURY phylum, TACK, DPANN, and Asgard superphyla within the Archaea domain. RecA is used as an outgroup in these trees. **(B).** A maximum-likelihood phylogenetic tree of Asgard RadA sequences, with E. coli RecA used as the outgroup. The branch confidence was determined by bootstrap re-sampling using ultrafast bootstrapping. **(C).** Block alignment protein sequences from 14 groups identified recombinases from Asgard Archaea. The length of each rectangular block reflects the overall length of the sequence conservation logo generated from a multiple sequence alignment of all unabridged sequences in each group. Each block associated with an Asgard group was aligned by the location of the Walker A box in the RecA domain. Also highlighted are the relative locations of the Walker B motif and the DNA binding loops. Also illustrated are N-terminal and C-terminal extensions unique to different groups of recombinase proteins.

### Extended View Figure 4

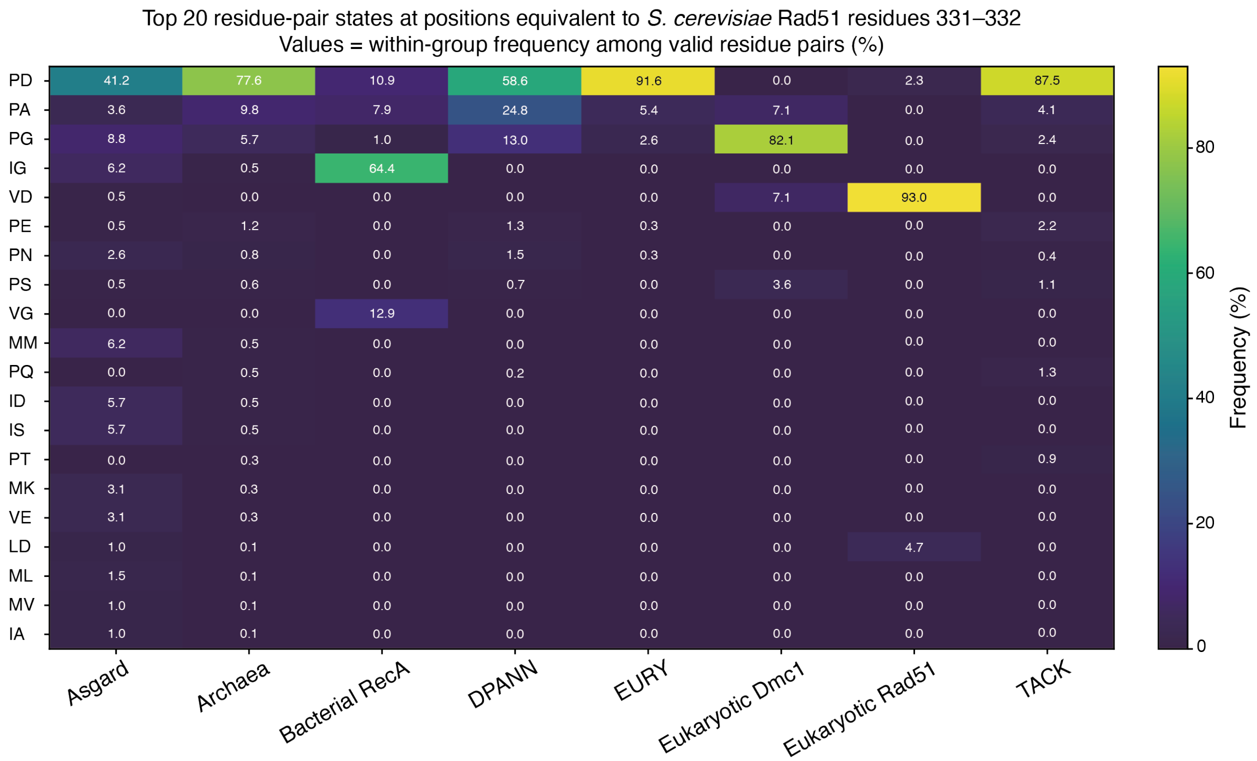


**EVFig 4: Distribution of common Loop 2 residue-pair states across recombinase groups**. The heatmap shows the frequencies of the 20 most abundant amino acid pair states at positions equivalent to *S. cerevisiae* Rad51 residues 331–332 across archaeal RadA subgroups, combined Archaea, bacterial RecA, and eukaryotic Rad51 and Dmc1 sequences. Pair states were ranked according to their frequencies in the combined dataset. Values indicate the within-group percentage among sequences containing non-gap amino acids at both positions.

### Extended View Figure 5

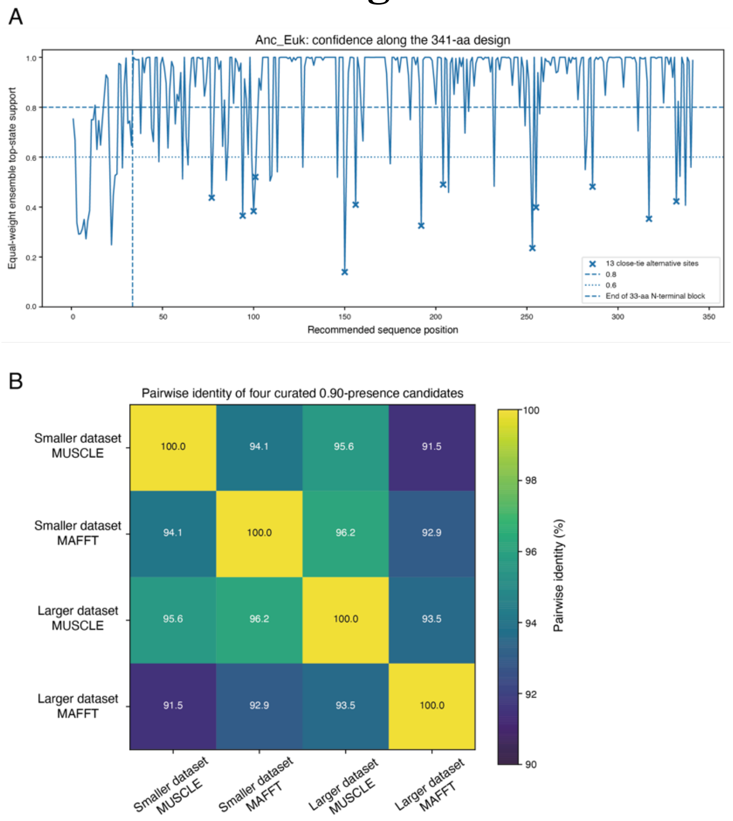


**EVFig 5:** Sequence-level support and cross-pipeline consistency of the recommended ancestral eukaryotic recombinase sequence.

**(A).** Sitewise ensemble support across positions of the 341-aa recommended Anc_Euk recombinase sequence. Amino acid posterior distributions from the available curated reconstructions were mapped to a common coordinate framework anchored to human RAD51. At each human-RAD51-anchored coordinate, posterior probabilities for each of the 20 amino acids were averaged with equal weight across all curated ASR pipelines in which that coordinate was represented. The curve shows the support for the top-ranked ensemble state. Horizontal lines indicate ensemble-support thresholds of 0.8 and 0.6. Crosses mark 13 close-tie core positions at which top-state support was below 0.8 and the support difference between the top- and second-ranked states was less than 0.15. The vertical dashed line marks the end of the alignment-sensitive, acidic/Q-rich 33-aa N-terminal block. The recommended N-terminal sequence was retained as a coherent block from the larger-dataset MUSCLE v5.1 reconstruction, whereas the remaining sequence was selected using the equal-weight ensemble analysis. **(B).** Pairwise sequence identities among four exploratory full-length ancestral candidates generated from the four untrimmed analyses of the curated smaller and larger datasets using the corresponding MUSCLE and MAFFT G-INS-I alignments. For each analysis, alignment columns with inferred $P(present) \geq 0.90$ were retained, and the maximum-posterior amino acid was selected at each retained position to generate a candidate sequence. The four candidate sequences were subsequently aligned, and pairwise identity was calculated over positions containing a non-gap amino acid in both sequences. Values within the heatmap indicate pairwise sequence identity (%).

### Extended View Figure 6

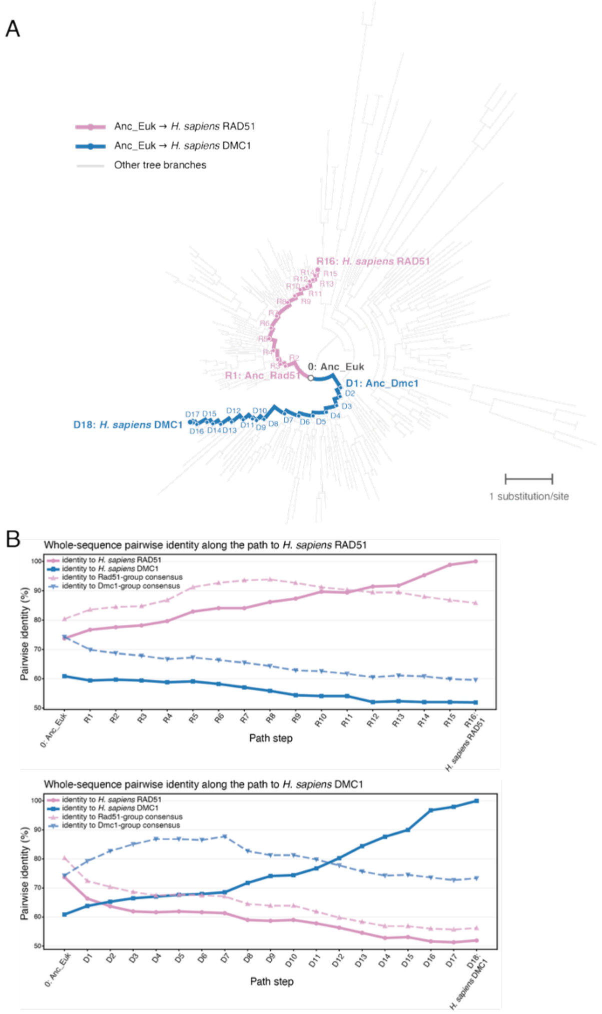


**EVFig6:** Comparison of evolutionary trajectories from Anc_Euk to *H. sapiens* RAD51 and DMC1.

**(A)**. Maximum-likelihood phylogeny generated from the full, non-curated RadA/Rad51/Dmc1 dataset described in Methods. The ancestral paths from Anc_Euk to *H. sapiens* RAD51 and *H. sapiens* DMC1 are highlighted in pink and blue, respectively, whereas all other branches are shown in light gray. R1–R16 and D1–D18 denote successive nodes along the RAD51 and DMC1 paths, respectively; the reconstructed Anc_Rad51 and Anc_Dmc1 nodes are indicated. Branch lengths represent substitutions per site. **(B)**. Whole-sequence pairwise identities of successive reconstructed nodes along the ancestral paths to *H. sapiens* RAD51 (top) and *H. sapiens* DMC1 (bottom). Pairwise identity was calculated across mutually comparable alignment positions containing a standard amino acid in both sequences; positions containing a gap or an ambiguous or non-standard residue in either sequence was excluded. At each path step, sequence identity was calculated relative to the extant human RAD51 and DMC1 sequences and to the consensus sequences of the Rad51 and Dmc1 groups. Solid lines with circles indicate identities to the extant human proteins, whereas dashed lines with triangles indicate identities to the Rad51 and Dmc1 group consensuses.

### Extended View Figure 7

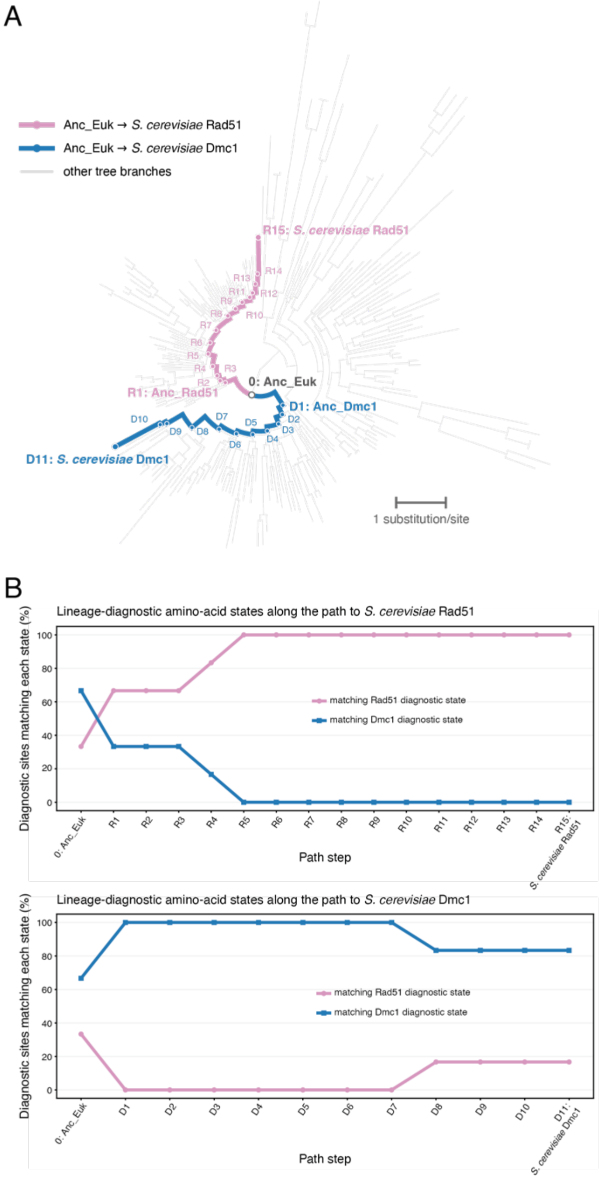


**EVFig 7: Comparison of evolutionary trajectories from Anc_Euk to *S. cerevisiae* Rad51 and Dmc1.**

**(A).** Maximum-likelihood phylogeny generated from the full, non-curated RadA/Rad51/Dmc1 dataset described in Methods. The ancestral paths from Anc_Euk to *S. cerevisiae* Rad51 and *S. cerevisiae* Dmc1 are highlighted in pink and blue, respectively, whereas all other branches are shown in light gray. R1–R15 and D1–D11 denote successive nodes along the Rad51 and Dmc1 paths, respectively; the reconstructed Anc_Rad51 and Anc_Dmc1 nodes are indicated. Branch lengths represent substitutions per site. **(B).** Changes in lineage-diagnostic amino acid states along successive reconstructed nodes on the ancestral paths to *S. cerevisiae* Rad51 (top) and *S. cerevisiae* Dmc1 (bottom). At each path step, the percentage of the six diagnostic positions whose reconstructed residues matched the Rad51-diagnostic or Dmc1-diagnostic state was calculated. The extant *S. cerevisiae* sequence is included as the final step along each path.

### Extended View Figure 8

A

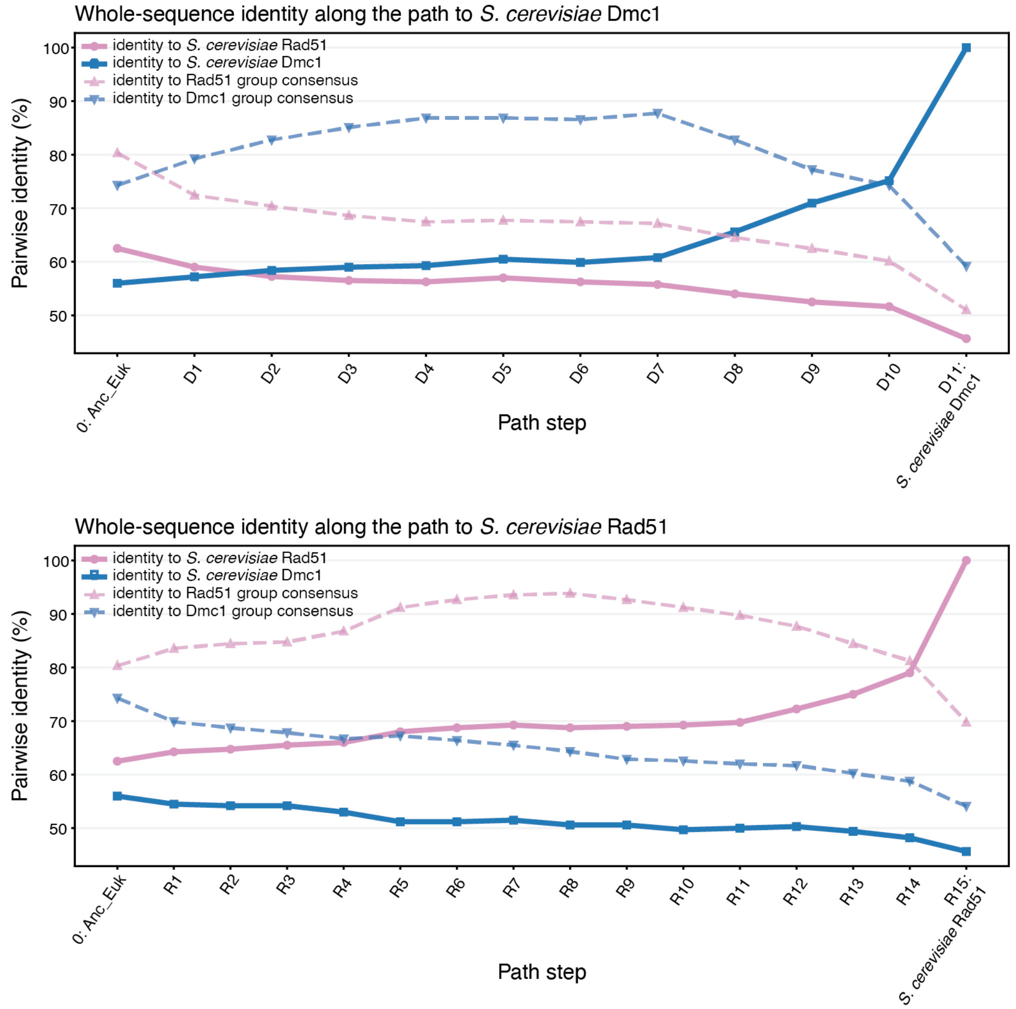


**EVFig 8:** Comparison of evolutionary trajectories from Anc_Euk to *S. cerevisiae* Rad51 and Dmc1 (whole-sequence pairwise identities).

**(A).** Whole-sequence pairwise identities of successive reconstructed nodes along the ancestral paths to *S. cerevisiae* Rad51 (top) and *S. cerevisiae* Dmc1 (bottom). Pairwise identity was calculated across mutually comparable alignment positions containing a standard amino acid in both sequences; positions containing a gap or an ambiguous or non-standard residue in either sequence was excluded. At each path step, sequence identity was calculated relative to the extant *S. cerevisiae* Rad51 and Dmc1 sequences and to the consensus sequences of the Rad51 and Dmc1 groups. Solid lines with circles indicate identities to the extant budding yeast proteins, whereas dashed lines with triangles indicate identities to the Rad51 and Dmc1 group consensuses.

### Extended View Figure 9

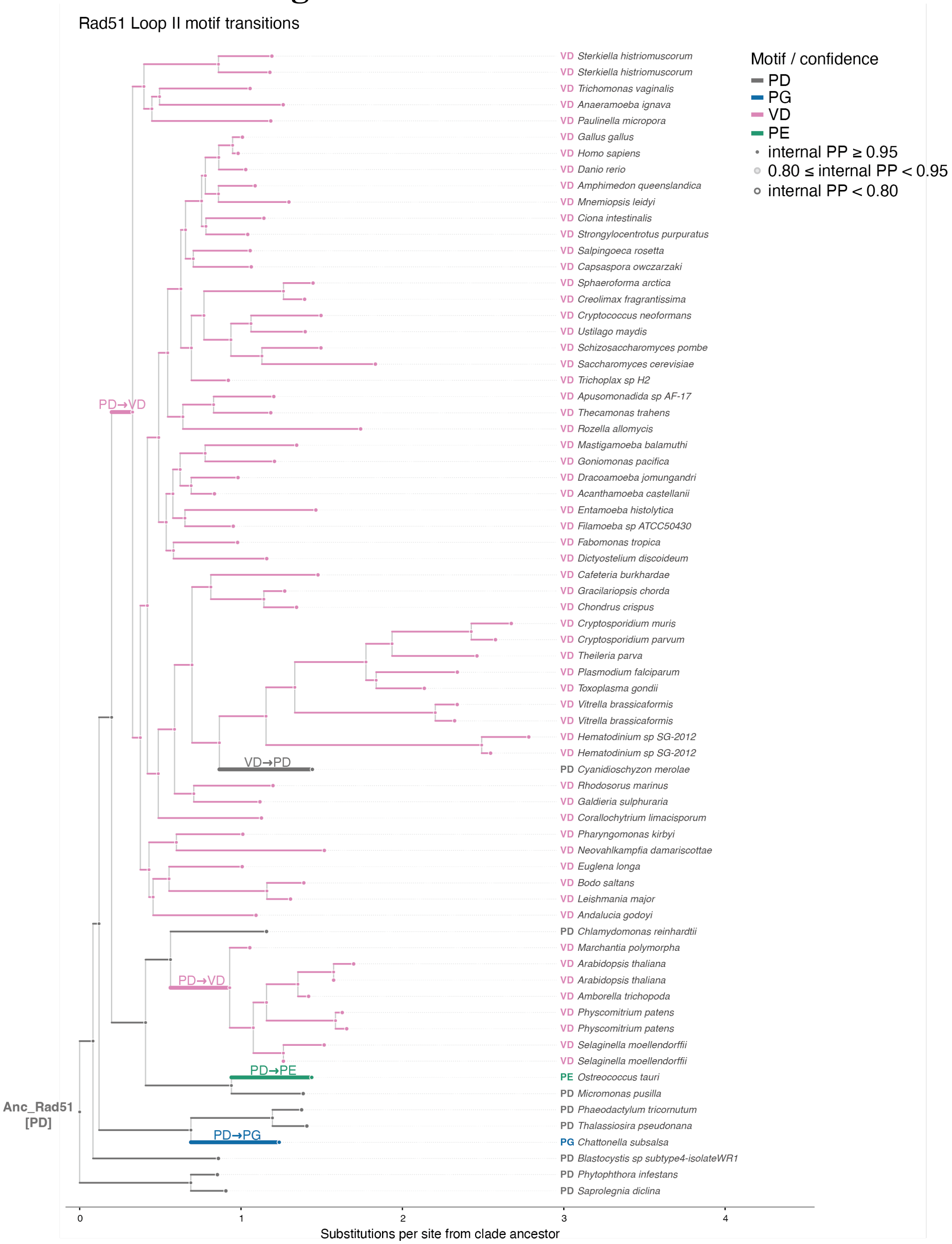

**EVFig 9: Detailed phylogenetic reconstruction of Loop 2 residue-pair state transitions in the Rad51 lineage.**

The Rad51 subtree was extracted from the maximum-likelihood phylogeny generated using the full, non-curated RadA/Rad51/Dmc1 dataset described in Methods. The reconstructed Anc_Rad51 state at positions equivalent to *S. cerevisiae* Rad51 residues 331–332 was PD. Branches are colored according to the reconstructed amino acid pair state, whereas terminal labels indicate the observed states of extant sequences. Labels along branches indicate inferred transitions between pair states. Node symbols indicate confidence in the reconstructed pair state, based on the lower of the two site-specific posterior probabilities (PPs), with confidence categories defined in the legend. Horizontal distances represent cumulative substitutions per site from Anc_Rad51; vertical spacing is for visualization only. This tree provides the detailed phylogenetic representation underlying the simplified transition map in Figure 2D.

### Extended View Figure 10

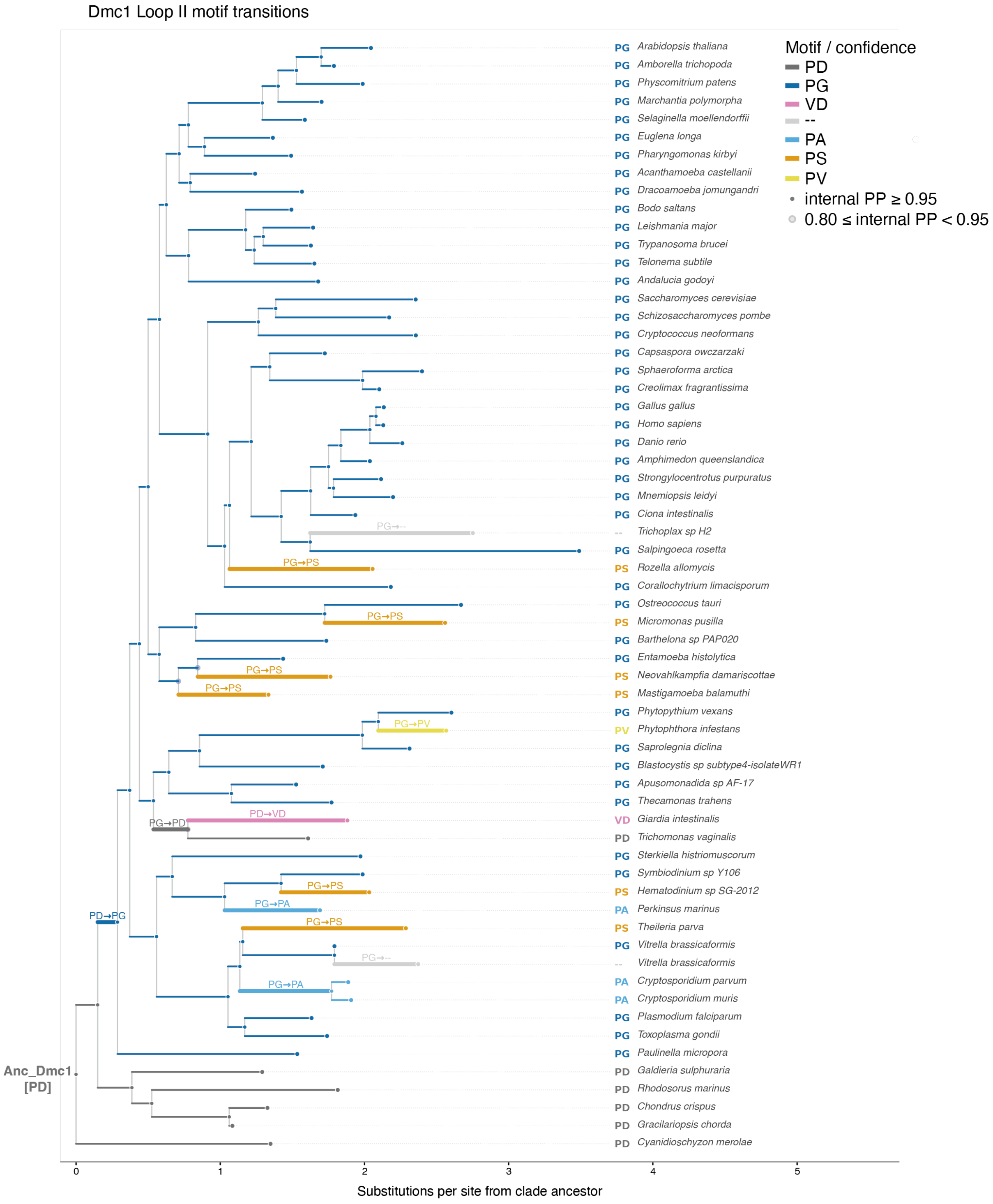

**EV Fig 10: Detailed phylogenetic reconstruction of Loop II residue-pair state transitions in the Dmc1 lineage.**

The Dmc1 subtree was extracted from the maximum-likelihood phylogeny generated using the full, non-curated RadA/Rad51/Dmc1 dataset described in Methods. The reconstructed Anc_Dmc1 state at positions equivalent to *S. cerevisiae* Rad51 residues 331–332 was PD. Branches are colored according to the reconstructed amino acid pair state, whereas terminal labels indicate the observed states of extant sequences. Labels along branches indicate inferred transitions between pair states. Node symbols indicate confidence in the reconstructed pair state, based on the lower of the two site-specific posterior probabilities (PPs), with confidence categories defined in the legend. Horizontal distances represent cumulative substitutions per site from Anc_Dmc1; vertical spacing is for visualization only. This tree provides the detailed phylogenetic representation underlying the simplified transition map in Figure 2E.
